# NFATC2 in pancreatic cancer-associated fibroblasts predicts treatment response and facilitates ERBB-targeted therapies

**DOI:** 10.64898/2026.04.04.716465

**Authors:** Jiahao Guo, Samuele Cancellieri, Chi Xu, Celina Wiik, Liangru Fei, Shiva Dahal-Koirala, Emma Haapaniemi, Tero Aittokallio, Caroline Sophie Verbeke, Biswajyoti Sahu

## Abstract

Pancreatic ductal adenocarcinoma (PDAC) remains a lethal malignancy, with therapeutic resistance influenced by a dense desmoplastic stroma dominated by cancer-associated fibroblasts (CAF). Using single-cell RNA-sequencing and gene regulatory network modeling of 42 PDAC tumors, we identified a CAF subpopulation characterized by elevated NFATC2 expression that is enriched in patients with improved therapeutic response and survival. NFATC2+ CAFs exhibited tumor-suppressive features, including enhanced apoptotic signaling and suppression of ERBB pathway activity. Co-culture experiments demonstrated that NFATC2+ CAFs restrain pancreatic cancer cell growth and enhance chemotherapy-induced apoptosis, increasing sensitivity to standard-of-care chemotherapy regimens and synergizing with ERBB-targeted therapies. The favorable effect of NFATC2+ CAFs on chemotherapy response was validated in two other PDAC cohorts and in rectal cancer. Together, these findings identify NFATC2+ CAFs as a therapy-conditioned stromal state linked to improved treatment response and uncover a context-dependent vulnerability within the tumor microenvironment that may be exploited to rationally optimize combination therapies.

## INTRODUCTION

Pancreatic ductal adenocarcinoma (PDAC) remains one of the most treatment-refractory human malignancies^1,2^, its dismal outcome reflecting aggressive tumor biology, lack of effective early detection strategies, and profound resistance to existing therapies^3–5^. Despite advances in surgery and systemic therapies including multi-drug chemotherapy regimens such as FOLFIRINOX^3,6^, median overall survival for most patients remains less than one year, underscoring a critical unmet need for strategies that overcome therapeutic resistance. A defining feature of PDAC is its exceptionally dense, desmoplastic tumor microenvironment (TME), which not only limits drug delivery but also actively shapes tumor cell adaptation under therapeutic pressure^5,7–9^. Notably, there are currently no validated molecular biomarkers to stratify patients for existing therapies or guide rational combination approaches, highlighting the need to better understand stromal determinants of treatment response in PDAC.

The desmoplastic stromal characteristic of PDAC can comprise up to 90% of the total tumor mass^10–12^. Cancer-associated fibroblasts (CAF) are the predominant cellular population in stroma that play pivotal roles in tumor progression, metastasis, and therapeutic resistance^13–15^. CAFs create a tumor-supportive microenvironment through growth factor secretion, extracellular matrix (ECM) remodelling, angiogenesis facilitation, metabolic crosstalk and immune modulation. Because CAFs dominate the PDAC tumor mass and actively constrain drug delivery and immune infiltration, therapeutic responses in PDAC are inherently shaped by stromal biology. Thus, CAF-directed approaches may be particularly impactful in complementing therapies targeting malignant cells^16^, and defining treatment-responsive CAF populations and their molecular regulators may reveal new targets for combination therapies that enhance treatment efficacy and overcome resistance.

Single-cell RNA sequencing (scRNA-seq) analyses have revealed substantial CAF heterogeneity within the PDAC TME^17–23^ and, importantly, mapped the stromal transcriptional responses to chemotherapy^24^. Recent evidence highlights transcriptional adaptation as a major determinant of both cancer progression^25^ and treatment outcome^26^. However, which CAF states actively contribute to therapy response and the transcriptional regulators that orchestrate CAF state transitions during PDAC progression remain largely unknown.

Transcriptional adaptation and cell-state transitions are controlled by transcription factors (TF), but their functional activity in the PDAC stroma is embedded within complex, dynamic regulatory networks that are not readily captured by gene expression alone. Gene regulatory network (GRN) approaches provide a powerful framework to infer TF activity and identify regulators of cell-state transitions^27^. While recent studies have linked stromal features to PDAC patient outcome and treatment response using machine-learning analyses of bulk RNA sequencing and histopathological images^28–30^, GRN-methods have not yet been applied to interrogate therapy-induced stromal reprogramming at single-cell resolution, leaving TF-level mechanisms of CAF plasticity largely unexplored.

Here, using an integrated GRN and single-cell framework, we identify NFATC2 as a transcriptional regulator defining a treatment-responsive CAF state in PDAC. Our precision oncology pipeline integrates GRN inference and scRNA-seq analysis of treatment-naïve and neoadjuvantly treated PDAC tumors, followed by functional validation in human CAF models. NFATC2, a calcium-responsive TF of the nuclear factor of activated T cells (NFAT) family, was selectively expressed in a distinct subset of CAFs associated with improved treatment response and patient survival. The NFAT family TFs are best characterized for their roles in T cells, but have also been shown to regulate for example cell proliferation, differentiation, and apoptosis in non-immune cells, exerting either tumor suppressive or oncogenic roles depending on the cancer type and cellular context ^31–33^. However, the specific role of NFATC2 in CAFs, and its contribution to PDAC progression and therapeutic response, remains largely unexplored.

Our results show that inducing NFATC2 expression in human CAFs impairs the growth of PDAC cells in co-culture systems and increases their sensitivity to chemotherapy and synergizes with ERBB-targeted inhibition. Together, these findings identify NFATC2 as a treatment-inducible regulator of CAF biology in PDAC and imply a stromal mechanism linked to therapeutic benefit. By defining a TF-driven CAF state associated with improved clinical outcomes, this study provides mechanistic insight into therapy-induced stromal reprogramming and highlights a targetable vulnerability within the PDAC TME.

## RESULTS

### Gene regulatory network inference identifies NFATC2 as a treatment-responsive transcription factor in PDAC

To implement a precision oncology platform for identifying key TFs mediating transcriptional adaptation during PDAC treatment, we integrated GRN inference analysis with functional validation using co-culture systems (Figure 1A). We analyzed single-nuclei RNA sequencing data from primary tumor samples of 42 PDAC patients, comprising a total of 224,988 cells^34^. Cells were classified into 20 major cell types (Figure 1B) and annotated according to patient treatment response based on the original dataset definitions (Figure 1C).

**Figure 1.**
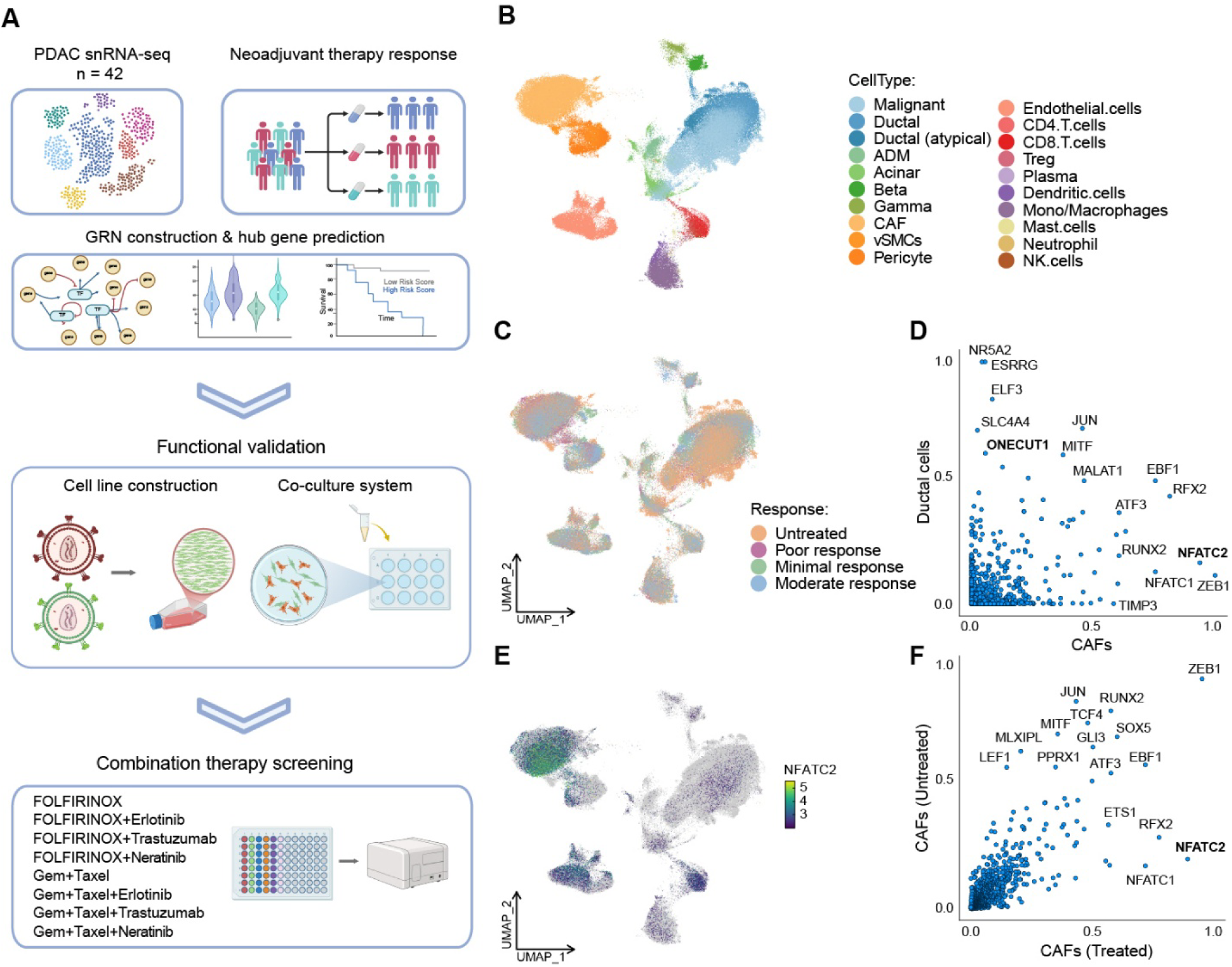
GRN construction reveals NFATC2 as a treatment-responsive TF in PDAC TME. (A) Workflow of our precision oncology pipeline integrating GRN construction and functional validation. (B) UMAP plot of 224,988 cells from 42 PDAC patients^34^ showing 20 cell types based on conventional markers (see Methods for details). (C) UMAP plot colored by treatment response categories as defined in the original dataset^34^, including tumors from untreated patients as well as patients receiving neoadjuvant therapy, classified as poor, minimal, or moderate responders, with moderate response representing the most favorable outcome. (D) Comparison of eigenvector centrality scores between ductal epithelial cells and CAFs, highlighting ONECUT1 enriched in ductal cell group and NFATC2 enrichment in the CAF group. (E) UMAP plot showing NFATC2 expression levels across different cell clusters. Color gradient represents normalized NFATC2 expression (log-transformed, purple to green: low to high expression). (F) Eigenvector centrality comparison between the CAFs from treated and untreated patients, highlighting NFATC2 enrichment in the treated group.

As a proof of concept, we first applied GRN inference using CellOracle^35^ to identify cell identity-specific TFs in pancreatic epithelial cell types. Eigenvector centrality analysis (a measure of node importance within regulatory networks that highlights putative master TFs) revealed several cell type-specific TFs, including ONECUT1 (HNF6) and ESRRG in ductal cells, MECOM in acinar cells, and NR5A2 in both populations (Supplementary Figure 1A), consistent with enriched expression of ONECUT1 and MECOM in ductal and acinar populations, respectively (Supplementary Figure 1B). These findings align with established roles of these TFs in pancreatic lineage specification: we previously identified ONECUT1 as a master TF in direct conversion of human fibroblasts into pancreatic ductal-like epithelial cells^36,37^, whereas MECOM is a known regulator of acinar-to-ductal lineage transitions^38,39^. To test the accuracy of the CellOracle predictions, we performed *in-silico* perturbation of MECOM, which induced a clear transition of acinar cells towards ductal identity when compared to random perturbation controls (Supplementary Figure 1C), reinforcing the validity of the inferred GRN.

Having validated GRN performance in epithelial lineages, we next applied this framework to identify key transcriptional regulators in stromal populations and to map their enrichment during chemotherapy treatment. GRN analysis between the ductal epithelial cells and the CAFs showed ductal cells being enriched for known pancreatic cell-specific TFs such as ONECUT1, ESRRG and NR5A2^40,41^, whereas the CAFs were enriched for ZEB1 and NFATC2 (Figure 1D). While ZEB1 has been reported to be associated with epithelial to mesenchymal transition (EMT) in multiple cancers and is a known mesenchymal cell marker^42^, the enrichment of NFATC2, a T-cell-associated TF, was particularly interesting. UMAP analysis showed that NFATC2 was almost exclusively expressed in CAFs (Figure 1E). Importantly, NFATC2 had the highest eigenvector centrality score in CAFs among the treated patients compared to untreated ones (Figure 1F), suggesting that neoadjuvant treatment may lead to NFATC2 expression in CAFs, whereas no such difference was observed for ZEB1.

To further support our findings on the transcriptional heterogeneity across the TME, we performed TF activity analysis across all cell types using SCENIC^43^. Hierarchical clustering revealed five distinct transcriptional modules (M1-M5) characterized by differential pathway enrichments (Supplementary Figure 1D). In module 5, NFATC2 emerged as a key factor associated with metabolic remodeling, TGF-β suppression, and cellular plasticity, suggesting its role in transcriptional reprogramming across the tumor ecosystem. Collectively, these results demonstrate the strength of GRN approaches in studying the transcriptional determinants of distinct cell populations in PDAC tumors and highlight NFATC2 as a treatment-responsive transcription factor in PDAC TME.

### NFATC2 defined a treatment-responsive CAF state linked to TME remodeling and improved outcome in PDAC

To further characterize the role of NFATC2 in CAFs and its association with therapeutic response in PDAC, we focused on single-cell analysis of stromal populations. Based on established markers^7,17^, we identified four distinct stromal subpopulations: inflammatory CAFs (iCAFs), myofibroblastic CAFs (myCAFs), pericytes, and vascular smooth muscle cells (vSMC; Figure 2A & Supplementary Figure 2A). In addition, we identified a CAF subcluster characterized by strong upregulation of NFATC2 (Figure 2A). These NFATC2-positive CAFs (NFATC2+ CAFs) constituted a subset within the iCAF population, concomitantly showing downregulation of myCAF markers such as MYH11 and aSMA (ACTA2)^44,45^ (Figure 2A, Supplementary Figure 2A & B). Stratification of stromal cells by treatment response revealed higher NFATC2 expression in tumors from patients showing better responses (moderate and minimal response groups) compared to untreated patients and those with poor response (Figure 2B). Consistently, density mapping demonstrated enrichment of the NFATC2+ CAF subpopulation in responders, whereas this population was markedly reduced in poor responders and almost undetectable in tumors of untreated patients (Supplementary Figure 2C).

**Figure 2.**
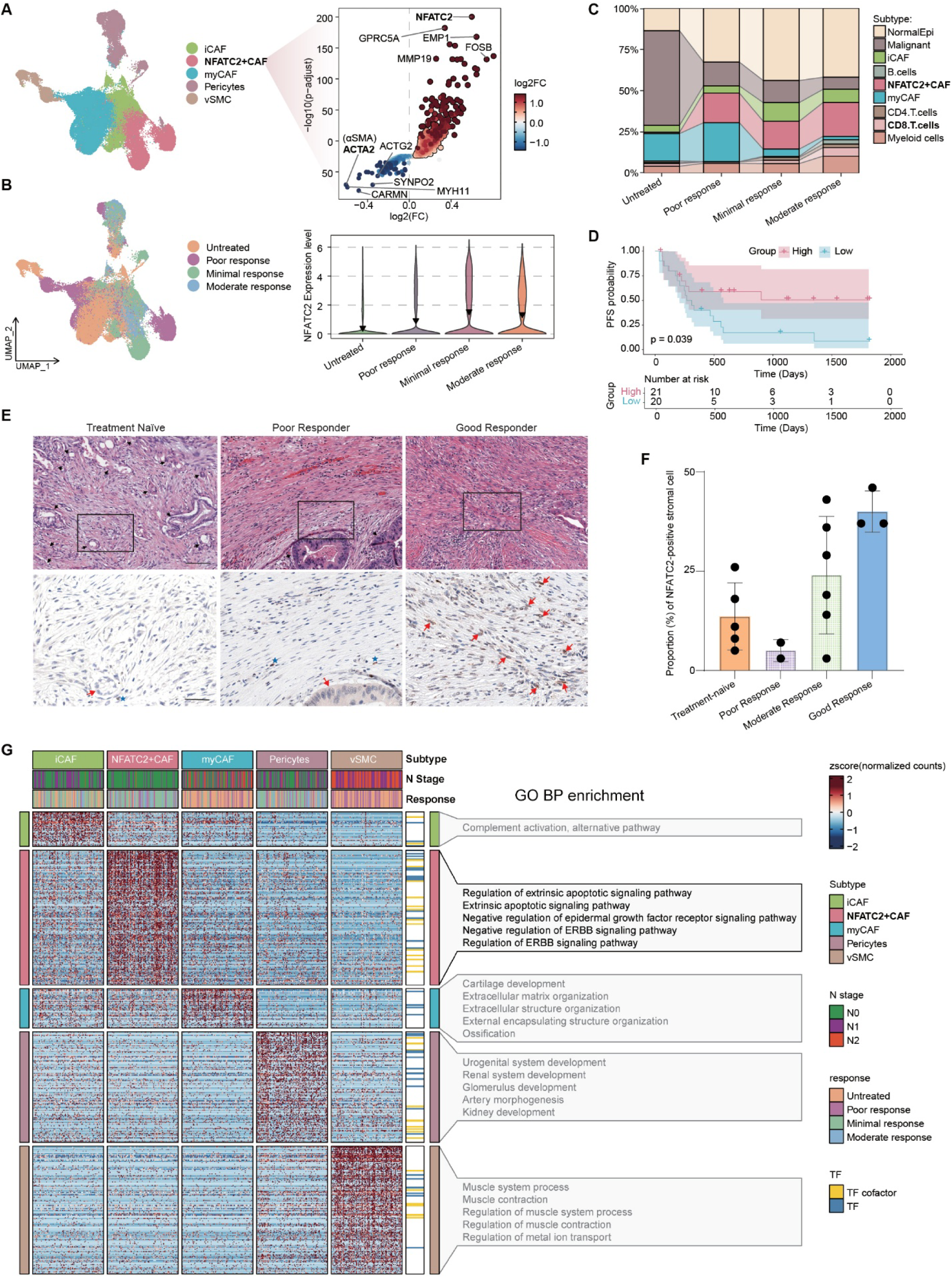
PDAC therapeutic response correlates with distinct CAF subpopulation distributions and NFATC2 expression levels. (A) Left: UMAP plots of stromal cells colored by subtype identity including inflammatory CAFs (iCAF), NFATC2-positive CAFs (NFATC2+ CAF), myofibroblastic CAFs (myCAF), pericytes, and vascular smooth muscle cells (vSMC). Right: Volcano plot for differential gene expression for NFATC2+ CAFs compared to other stromal cells. (B) Left: UMAP plots of stromal cells colored by patient therapy response categories (untreated, poor response, minimal response, and moderate response). Right: Violin plot showing NFATC2 expression levels across different treatment response groups. (C) Alluvial diagram quantifying the relative proportional distribution of cell types across different therapeutic response groups. Y-axis represents the percentage of cells (0-100%), with each cell type shown in distinct colors. The width of each bar reflects the relative sample size per response category. (D) Kaplan-Meier survival analysis demonstrates that patients with high NFATC2 (red line) based on the median pseudobulk expression in CAFs exhibit significantly longer progression-free survival compared to those with low expression (blue line, p = 0.039, log-rank test). The risk table shows patient numbers at different time points. (E) Representative images from tissue microarray analysis of PDAC patient samples stratified by treatment response. Upper panels show H&E staining of stromal regions from treatment-naive tumors (left), poor responder tumors (middle), and good responder tumors (right) following chemotherapy. Lower panels display corresponding NFATC2 immunohistochemistry staining of serial sections from the same regions. Black boxes indicate representative fields shown at higher magnification. NFATC2-positive CAFs (brown nuclear staining) are more abundant in good responder samples compared to poor responders and treatment-naive samples. Scale bars = 100 μm. (F) Quantification of NFATC2-positive CAF percentage across patient response groups. Bar graph shows percentage of NFATC2-positive stromal cells in treatment-naive tumors (orange, n=5), poor responders (pink, n=2), moderate responders (green, n=6), and good responders (blue, n=3). Each dot represents an individual patient sample. NFATC2 positivity progressively increases with better treatment response, with good responders showing markedly higher NFATC2+ CAF percentage (∼40%) compared to poor responders (∼5%) and treatment-naive samples (∼12%). Data are presented as mean ± SEM. (G) Heatmap displaying functional enrichment analysis results for each stromal cell subtype across different therapeutic response groups and N stage. Gene Ontology Biological Process (GO BP) terms are grouped by functional categories. The color scale represents normalized enrichment scores, with red and blue indicating positive and negative enrichment, respectively.

In parallel with increased abundance of NFATC2+ CAFs and higher NFATC2 expression levels in treatment responders, therapeutic response was further associated with shifts in the cellular composition of the TME. Tumors from the treatment-responding patients exhibited higher relative proportions of normal epithelial cells, iCAFs, CD8+ T cells, and myeloid cells compared to poor responders and untreated patients, whereas the relative proportions of malignant cells and myCAFs were reduced (Figure 2C). Together, these findings suggest a broad therapy-associated remodeling of the tumor ecosystem, rather than changes restricted to the CAF compartment, consistent with previous single-cell and spatial analyses of neoadjuvant-treated PDAC^34^.

NFATC2 expression in CAFs also correlated with favorable clinical features, including smaller tumor size and reduced lymph node involvement (lower T and N stages; Supplementary Figure 2D), and was highest in patients who had received neoadjuvant FOLFIRINOX-based chemoradiotherapy (CRT), compared to treatment-naïve patients and those receiving other neoadjuvant therapies (Supplementary Figure 2D). At the transcriptional level, the expression of marker genes associated with a recently described reactive subTME linked to chemosensitivity^20^ were increased in NFATC2+ CAFs compared to iCAF and myCAF populations, whereas the markers for deserted subTME were reduced (Supplementary Figure 2E). Importantly, stratification based on NFATC2 expression in CAFs revealed significantly prolonged progression-free survival in patients with high NFATC2 levels (Figure 2D), with concordant improvements in overall survival in the TCGA PDAC cohort (Supplementary Figure 2F), suggesting a protective, treatment-associated role for NFATC2 in PDAC.

To validate the clinical relevance of our findings in another patient cohort, we performed immunohistochemical analysis of NFATC2 expression in a tissue microarray comprising PDAC tumor samples with documented chemotherapy response. Consistent with our scRNA-seq observations, NFATC2+ CAFs were rare in treatment-naive tumors but became enriched in post-treatment samples (Figure 2E & Supplementary Table 1). Importantly, NFATC2 positivity in stromal compartments correlated strongly with treatment response, with good responders showing higher percentages of NFATC2+ CAFs (∼40%) compared to poor responders (∼5%) (Figure 2F). This clinical validation supports the translational relevance of NFATC2 as a stromal biomarker of treatment response in PDAC.

To understand the downstream signaling pathways associated with NFATC2+ CAFs, we performed Gene Ontology enrichment analysis for differentially expressed genes across all stromal cell types (Figure 2G & Supplementary Table 2). NFATC2+ CAFs showed enrichment for pathways related to regulation of extrinsic apoptotic signaling and negative regulation of ERBB signaling, whereas iCAFs were associated with complement activation. Other CAF types were enriched for pathways related to ECM organization and muscle system processes.

Collectively, these results identify NFATC2+ CAFs as a treatment-responsive stromal state in PDAC that is linked to improved patient survival and treatment outcome, enhanced apoptotic signaling, and suppression of ERBB pathways, suggesting that NFATC2 marks a stromal state associated with therapy-responsive remodeling of the PDAC TME.

### NFATC2 expression in CAFs limits PDAC cell growth and enhances apoptosis

To directly test the functional role of NFATC2 in CAFs, we established an optimized in vitro co-culture system using fluorescently labelled PANC1 pancreatic cancer cells expressing mCherry and human pancreatic CAFs (pCAF) expressing EGFP. Because pCAFs do not endogenously express NFATC2, we generated pCAFs with stable NFATC2 expression (NFATC2+ pCAFs; Figure 3A & Supplementary Figure 3A) and optimized CAF-to-cancer cell co-culture ratios based on previous reports^45–47^. pCAF and cancer cells cultured in a 3:1 ratio yielded stable co-culture conditions without either cell type outcompeting the other over time (Supplementary Figure 3B&C) and was used for all subsequent experiments.

**Figure 3.**
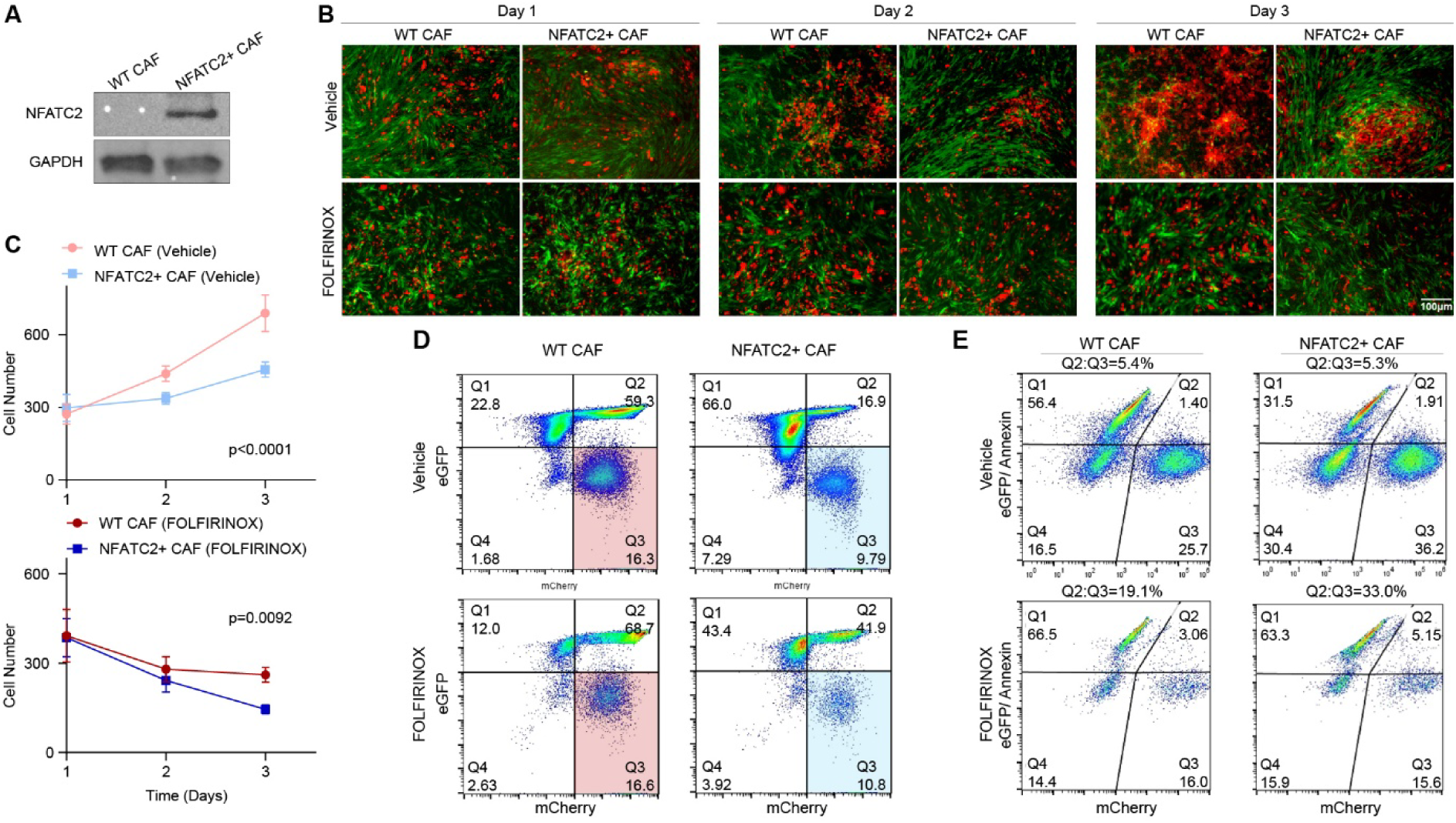
NFATC2 expression in CAFs limits PDAC cell growth and enhances apoptosis. (A) Western blot showing ectopic NFATC2 expression in pancreatic CAFs (pCAF) generated using lentiviral transduction. GAPDH served as loading control. WT, wild-type pCAFs; NFATC2+, NFATC2-expressing pCAFs. Uncropped blots are shown in Supplementary Figure 3A. (B) Representative fluorescence microscopy images of PANC1-pCAF co-cultures. PANC1 cells are labeled with mCherry (red) and pCAF cells with EGFP (green). WT and NFATC2+ pCAF co-cultures are shown on days 1-3 under vehicle or FOLFIRINOX treatment. Scale bar, 10μm. (C) Quantification of PANC1 cell numbers using CellProfiler from fluorescence images showing that NFATC2+ CAFs significantly reduce cancer cell proliferation, particularly under FOLFIRINOX treatment. Data are presented as mean ± SEM (n=3 independent experiments). Upper panel: p<0.0001. Lower panel: p=0.0092, two-way ANOVA. (D) Flow cytometry analysis of PANC1-pCAF co-culture systems with WT pCAFs (left) and NFATC2+ pCAF (right). Scatter plots display mCherry fluorescence (x-axis) versus eGFP fluorescence (y-axis), PANC1 cells (mCherry+, eGFP-) seen in the lower right quadrant. (E) Annexin V-FITC apoptosis assay by flow cytometry. Density plots showing the fraction of apoptotic cells (Q2, top-right quadrant) relative to live cancer cells (Q3, bottom-right quadrant) following co-culture with WT or NFATC2+ pCAFs ± FOLFIRINOX. In FOLFIRINOX-treated cells, the apoptotic fraction increased from 19.1% in WT pCAF co-cultures to 33.0% in NFATC2+ pCAF co-cultures, indicating enhanced chemotherapy-induced apoptosis in cancer cells co-cultured with NFATC2+ pCAFs.

PANC1 cells co-cultured with NFATC2+ pCAFs exhibited markedly altered growth dynamics compared to those co-cultured with wild-type (WT) CAFs: NFATC2+ pCAF co-cultures resulted in reduced cancer cell expansion within 48 hours (Figure 3B, top). Treatment of the co-cultures with the standard-of-care chemotherapy regimen FOLFIRINOX further strengthened this effect, resulting in a marked reduction in cancer cell numbers in NFATC2+ pCAF co-cultures compared to WT controls (Figure 3B, bottom). Quantification by fluorescence-based cell counting and flow cytometry confirmed that NFATC2 expression in CAFs limits cancer cell growth, and that FOLFIRINOX treatment further enhances this effect (Figure 3C&D). A similar response was observed in independent experiments using another pancreatic cancer cell line MIA PaCa-2 (Supplementary Figure 3D&E), supporting the generalizability of this effect.

Given that NFATC2+ CAFs were associated with apoptotic signaling in patient-derived single-cell analyses, we next assessed whether NFATC2 expression in CAFs promotes cancer cell apoptosis in the co-culture system. Flow cytometric analysis using Annexin V-FITC staining showed enhanced cancer cell apoptosis in NFATC2+ pCAF co-cultures, particularly under FOLFIRINOX treatment (Figure 3E). Together, these results establish that NFATC2 expression in CAFs attenuates pancreatic cancer cell growth, enhances chemotherapy-induced apoptosis, and sensitizes cancer cells to FOLFIRINOX.

### Transcriptional programs and paracrine effects underlying NFATC2-dependent CAF-cancer cell interactions

To define transcriptional programs underlying the NFATC2-dependent phenotypes observed in co-culture (Figure 3), we performed bulk RNA-seq under four complementary conditions: PANC1 cells co-cultured directly with WT or NFATC2+ pCAFs, or exposed to conditioned medium from WT or NFATC2+ pCAF cultures, each treated with either FOLFIRINOX or vehicle. Differential gene expression for each condition was analyzed between NFATC2+ and WT samples (Supplementary Table 3-6).

Pathway-level analysis using Gene Set Variation Analysis (GSVA) revealed robust activation of apoptosis-related programs, including Hallmark Apoptosis and p53 pathways in FOLFIRINOX-treated PANC1 cells exposed to NFATC2+ CAF-conditioned medium (Figure 4A), consistent with enhanced chemotherapy-induced DNA damage response. Both intrinsic (mitochondrial) and extrinsic (death receptor) apoptotic pathways were induced by NFATC2+ CAFs, with extrinsic signaling further enhanced by FOLFIRINOX (Figure 4B), consistent with increased measured apoptosis.

**Figure 4.**
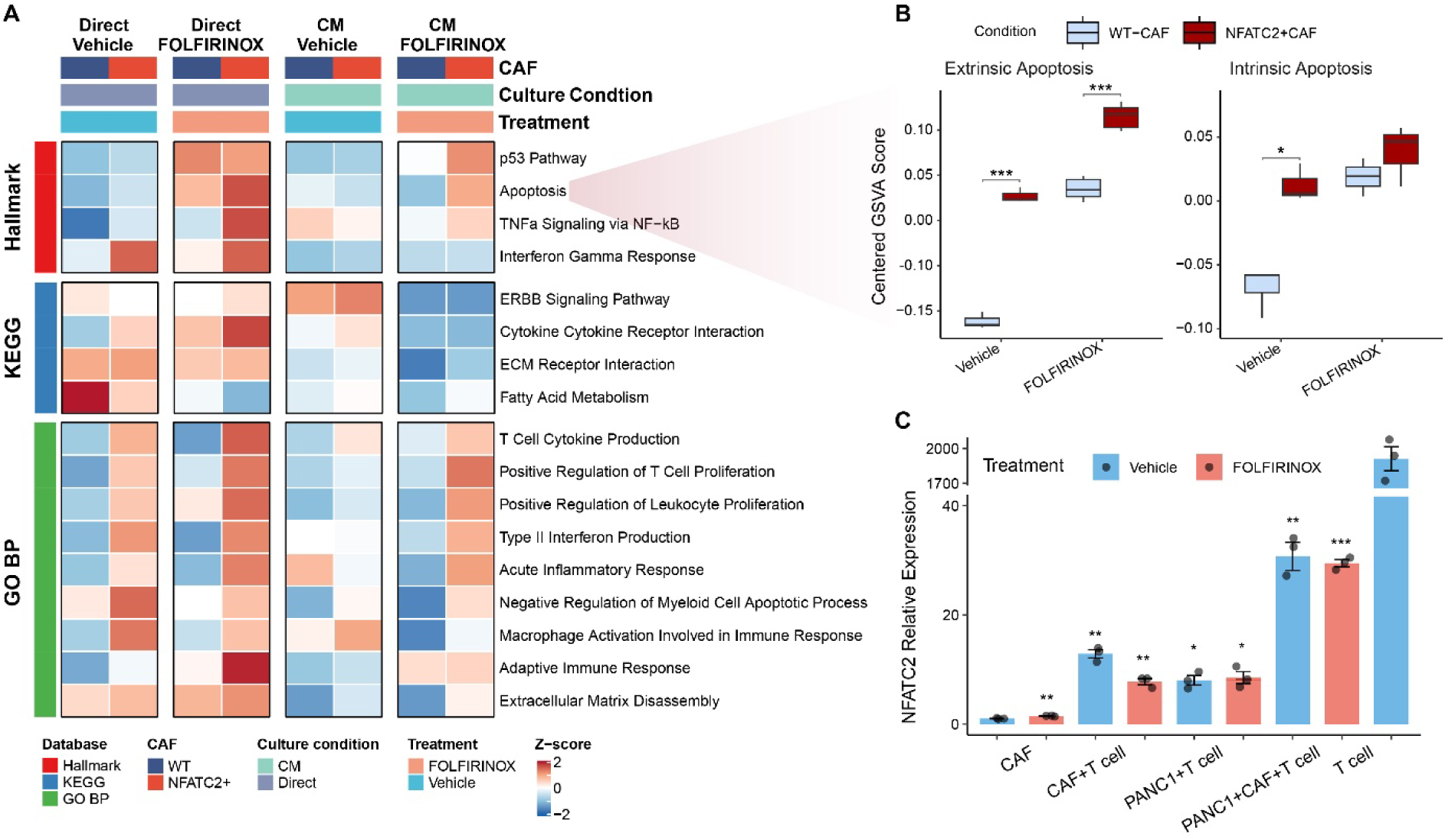
Transcriptional programs and paracrine effects underlying NFATC2-dependent CAF-cancer cell interactions. (A) GSVA heatmap showing pathway activity differences between WT and NFATC2+ CAF-conditioned PANC-1 cells across four experimental conditions (direct co-culture and conditioned medium, each with vehicle or FOLFIRINOX treatment). Each condition contains paired samples (WT and NFATC2+, representing merged biological triplicates). GSVA enrichment scores were calculated for each sample and averaged across biological triplicates. Rows represent selected pathways from Hallmark, KEGG and GO BP databases. Color intensity indicates GSVA enrichment Z-score (red, upregulated; blue, downregulated). Left annotation bar indicates pathway database source, top annotation bars CAF genotype (WT, blue; NFATC2+, red), culture method (direct, blue; CM, teal), and treatment (vehicle, cyan; FOLFIRINOX, orange). (B) GSVA enrichment analysis of apoptotic pathways from RNA-seq data. Box plots comparing extrinsic (left) and intrinsic (right) apoptosis pathway scores between WT CAF and NFATC2+ pCAF co-cultures ± FOLFIRINOX. ***p < 0.001, *p < 0.05 (t-test). (C) NFATC2 expression levels in different cell populations after 48-hour co-culture with activated T cells under vehicle (blue) and FOLFIRINOX (red) treatments. Five experimental groups were analyzed: (1) WT CAFs cultured alone; (2) WT CAFs co-cultured with T cells; (3) PANC-1 cells co-cultured with T cells; (4) PANC-1 and WT CAF co-culture system stimulated with T cells (PANC-1+ CAF+ T cell); (5) T cells cultured alone. Of note, T cells were removed from the co-cultures by washing before RNA extraction and qPCR analysis, ensuring that measured NFATC2 expression represents only CAF or PANC-1 cells. Data are presented as mean ± SEM of NFATC2 expression relative to vehicle CAF (n=3 biological replicates per group). Statistical significance was determined by unpaired Student’s t-test comparing vehicle vs FOLFIRINOX within each cell type. P-values compared with CAF Vehicle group are indicated above bars. ***p < 0.001, **p < 0.01, *p < 0.05 (t-test).

In direct CAF-PANC1 co-cultures, NFATC2 expression in CAFs was associated with prominent activation of inflammatory pathways and immune response programs (Figure 4A). Upregulation of inflammatory chemokines and cytokines (e.g. CXCL2/3/8, IL1A, IL1B, IL33) and immune signaling molecules (TREM1, TNFSF9/15, PTGS2), together with downregulation of myCAF-associated genes (e.g. CXCL12, COL11A1, MFAP5, MYLK) indicated NFATC2-dependent stromal reprogramming towards an iCAF phenotype (Figure 4A & Supplementary Figure 4A & Supplementary Table 3&4). Interestingly, genes linked to Ca^2+^-signaling, such as GRIA1, were also up-regulated in NFATC2+ conditions (Supplementary Figure 4A), suggesting a potential mechanism for calcium-dependent NFATC2 activation^48^. FOLFIRINOX further reinforced inflammatory and stress-adaptive programs and was associated with enrichment of immune response and T cell-related pathways, suggesting a more immune-permissive stromal environment. Concordantly, NFATC2 expression correlated with reduced CXCL12 levels (Supplementary Figure 4A), a stromal feature that has previously been linked to enhanced T-cell infiltration in pre-clinical PDAC models^49^

Conditioned medium from NFATC2+ CAFs exerted a strong transcriptional effect on PANC1 cells, particularly in the presence of FOLFIRINOX (Supplementary Table 5&6), indicating that NFATC2+ CAFs modulate tumor cell responses, at least in part, through paracrine mechanisms. Notably, ERBB signaling was strongly suppressed in FOLFIRINOX-treated, conditioned-medium-exposed PANC1 cells (Figure 4A), consistent with ERBB pathway downregulation observed in the patient-derived scRNA-seq data (c.f. Figure 2F). Notably, the cytokine storm observed in direct co-cultures was absent in conditioned-medium–treated PANC1 cells, indicating that inflammatory signals predominantly originate from the CAF compartment.

Analysis of ECM programs revealed context-dependent NFATC2 effects. In direct CAF–PANC1 co-cultures, NFATC2 expression was associated with attenuation of myCAF programs and induction of dynamic, wound-healing–associated remodeling, marked by COL15A1, TNFAIP6, SERPINE2 and matrix-modifying enzymes such as ADAMTS5 and CTSK (Supplementary Figure 4A). In contrast, conditioned-medium-treated PANC1 cells showed a tumor-intrinsic response characterized by epithelial stabilization and anchoring to the basement membrane, with upregulation of structural ECM scaffolding components (HSPG2, LAMA5, COL18A1, COL8A2, COL5A3) and adhesion molecules, including factors maintaining epithelial architecture such as SPINT2^50^ (Supplementary Figure 4B). This basement-membrane-associated epithelial containment, most pronounced under FOLFIRINOX treatment, provides a potential mechanistic link between NFATC2 expression and improved therapeutic response.

To explore potential signaling mechanisms underlying these paracrine effects, we performed ligand–receptor interaction analysis^51^ using scRNA-seq data across CAF subtypes. NFATC2+ CAFs exhibited distinct secretory profiles, including elevated expression of candidate factors such as MMP14 and nicotinamide phosphoribosyltransferase (NAMPT) (Supplementary Figure 4C), suggesting potential routes of CAF–cancer cell communication.

Having established broad functional and transcriptional effects of NFATC2+ CAFs on cancer cells, we next investigated how this CAF state could arise within the TME. Because NFATC2+ CAFs were largely absent in untreated condition but enriched following therapy, we asked whether treatment-associated microenvironmental cues could induce NFATC2 expression in otherwise NFATC2-negative CAFs. Given the well-established role of NFATC2 in T cells, we examined whether immune cells could promote NFATC2 induction in pCAFs. While NFATC2 was undetectable in WT pCAFs cultured alone, robust induction was observed upon co-culture with T cells (Figure 4C), with the strongest upregulation observed in triple co-culture conditions involving WT pCAFs, T cells, and PANC1 cancer cells. Of note, these quantitative PCR measurements were performed after removing the T cells from the co-cultures, confirming that NFATC2 induction occurred within the CAF compartment. These findings suggest that immune-derived signals represent one potential trigger for the emergence of NFATC2+ CAF states within the TME, providing a potential link between treatment-associated immune activity and stromal reprogramming.

Collectively, these results position NFATC2 as a regulator of CAF plasticity, linking immune-mediated induction of NFATC2 to paracrine transcriptional programs that modulate cancer cell behavior and chemotherapy response.

### ERBB pathway and proliferative programs are suppressed in treatment-responsive cancer cells

The enrichment of extrinsic apoptosis pathway and negative regulation of ERBB signaling pathway in the NFATC2+ CAFs from better-responding patients and NFATC2+ CAF-PANC1 co-cultures (c.f. Figures 2F&4A) prompted us to examine transcriptional changes within the cancer cell population of these patients from the scRNA-seq data^34^. For this, we separated malignant cells from other epithelial cells based on their copy number variation profiles (Supplementary Figures 5A-C) and performed transcriptomic analyses of malignant cells stratified by neoadjuvant therapy response. This revealed distinct transcriptional states based on neoadjuvant therapy response (Figure 5A).

**Figure 5.**
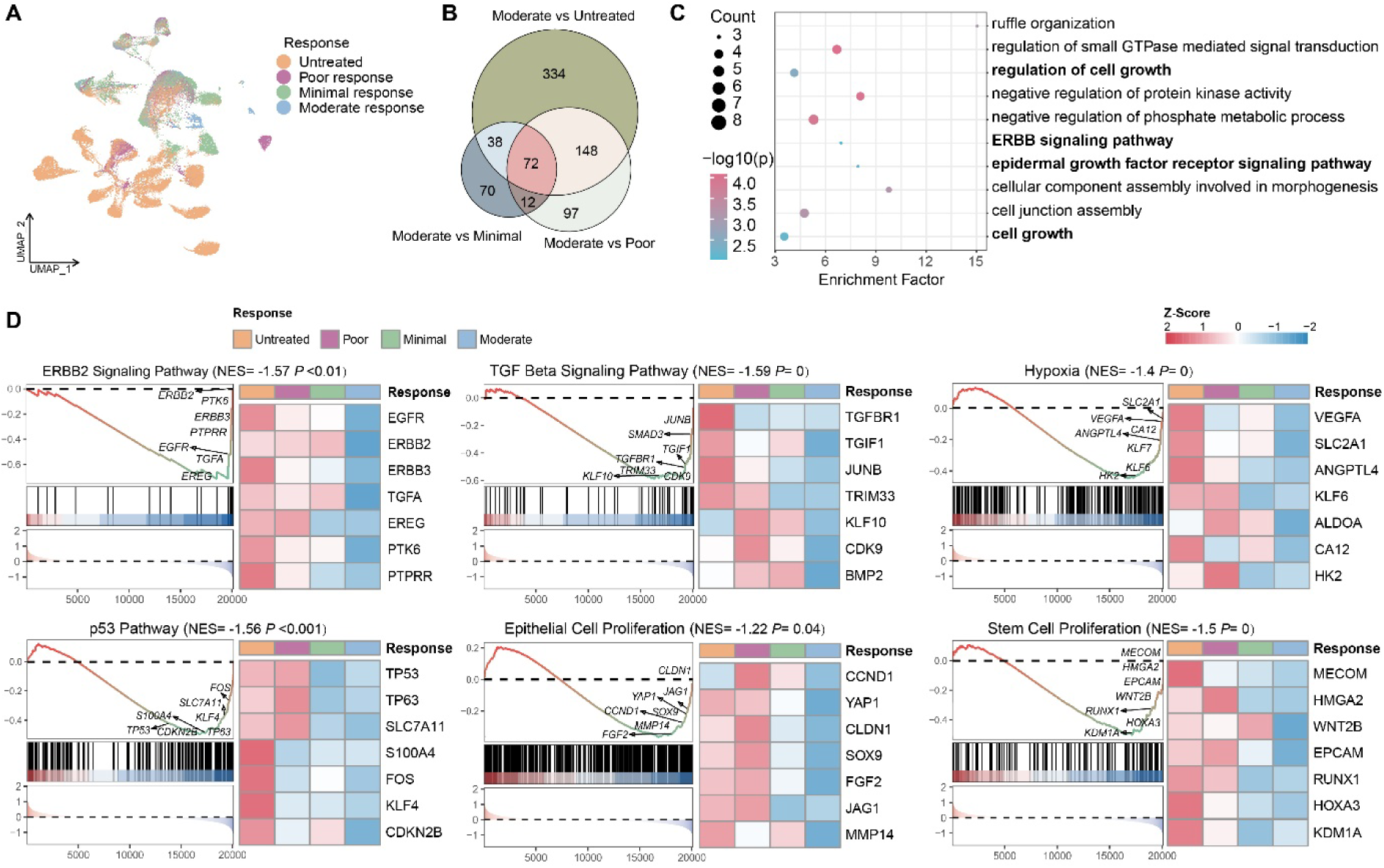
Transcriptional signatures of tumor cells stratified by NFATC2 expression in CAFs and treatment response. (A) UMAP visualization of single-cell transcriptomes of malignant cells from patients stratified by CAF NFATC2 expression levels and treatment response. Tumor cells are colored by patient treatment response classification: untreated (orange), poor response (purple), minimal response (green), and moderate response (blue). (B) Venn diagram illustrating the overlap of differentially expressed genes in tumor cells between response group comparisons. Numbers indicate unique and shared genes between Moderate vs Untreated (334 unique), Moderate vs Minimal (70 unique), and Moderate vs Poor (97 unique), with 72 shared genes commonly differentially expressed across all three comparisons and varying degrees of pairwise overlap. (C) Dot plot showing enriched biological processes and signaling pathways in the 72 overlapped genes from GO and KEGG pathway enrichment analysis. The x-axis represents the enrichment factor, the y-axis shows statistical significance as -log10(p-value), dot size indicates the number of genes in each category (3-8 genes), and dot color intensity represents the p-value. (D) Gene Set Enrichment Analysis (GSEA) plots of six cancer-relevant pathways in tumor cells comparing moderate responders to other response groups. Each panel displays the enrichment plot (left) with running enrichment score and rank metric, accompanied by a heatmap (right) showing normalized expression of leading-edge genes across response groups. Normalized negative enrichment scores (NES) indicate pathway downregulation in moderate response group compared to other responders. Analyzed pathways include ERBB2 Signaling Pathway (NES = -1.57, P = 0.01), TGF Beta Signaling Pathway (NES = -1.59, P < 0.0001), Hypoxia (NES = -1.4, P < 0.0001), p53 Pathway (NES = -1.56, P < 0.001), Epithelial Cell Proliferation (NES = -1.22, P = 0.04), and Stem Cell Proliferation (NES = -1.5, P < 0.001). Leading-edge genes driving pathway enrichment are labeled on the enrichment curves. Heatmap displays z-score normalized expression of those genes ranging from -2 (blue, low expression) to +2 (red, high expression).

Differential expression analysis identified genes associated with distinct treatment responses, with 72 genes consistently differentially expressed in moderate responders compared to each of the other response groups (Figure 5B). Gene Ontology enrichment analysis of the shared genes revealed significant enrichment of biological processes central for cancer cell growth and survival (Figure 5C), including regulation of cell proliferation, ERBB signaling, and epidermal growth factor receptor signaling pathways.

Gene Set Enrichment Analysis (GSEA) revealed significant negative enrichment of several cancer-relevant gene sets in treatment-responsive tumors (Figure 5D). Notably, ERBB2 signaling was markedly suppressed in moderate responders, consistent with reduced ERBB pathway activity observed in the stromal compartment of better-responding tumors enriched for NFATC2+ CAFs. In addition, malignant cells from moderate responders exhibited coordinated suppression of TGF-beta signaling, hypoxia response, p53 pathway activity, and proliferation programs, all of which are well-established drivers of tumor cell survival, plasticity, and therapy resistance^52^. These pathways are also central in CAF biology and tumor-stroma crosstalk^16^, providing an important link between transcriptional states in malignant cells and NFATC2-associated stromal environment. In particular, TGF-β signaling is a primary driver of myCAF differentiation and CAF activation across multiple cancer types^53^, while hypoxia, a characteristic feature of the PDAC TME, has been shown to promote iCAF phenotype through HIF-1α stabilization and cytokine-mediated paracrine signaling^54^.

Together, these results indicate that favorable therapeutic response in PDAC is associated with coordinated suppression of proliferative and ERBB signaling programs in cancer cells, highlighting ERBB pathway modulation as a potential vulnerability in treatment-responsive disease states.

### ERBB-targeted combination therapies enhance chemotherapy efficacy in NFATC2+ pCAF co-cultures

EGFR- and ERBB-targeted inhibitors have previously been tested in PDAC, but have shown no survival benefit in patients^55^ and only limited efficacy in pre-clinical models^56^. Our results from the PDAC patient data demonstrated that the better responding tumors are characterized by NFATC2+ pCAFs, suppression of ERBB signaling programs, and enhanced sensitivity to chemotherapy. This suggests that NFATC2 expression could mark treatment-responsive stromal contexts, potentially facilitating patient stratification for adjuvant or combination therapies. This prompted us to investigate whether ERBB-targeted therapies could potentiate standard-of-care chemotherapy in an NFATC2-positive stromal context.

For this, we performed a combinatorial drug testing using optimized PANC1-pCAF co-culture system comparing the effects in WT and NFATC2+ pCAFs conditions. Standard chemotherapy regimens (FOLFIRINOX or gemcitabine plus nab-paclitaxel) were administered alone or in combination with clinically relevant ERBB-targeted inhibitors including trastuzumab (anti-ERBB2 (HER2) monoclonal antibody), erlotinib (EGFR tyrosine kinase inhibitor), and neratinib (pan-ERBB inhibitor). Fluorescence microscopy analysis under vehicle-treated conditions confirmed the growth-suppressive effect of NFATC2 expression in CAFs on PANC1 cells (c.f. Figure 3). While FOLFIRINOX alone reduced cancer cell numbers, the addition of ERBB-targeting agents produced significantly greater cytotoxic effects, particularly in NFATC2+ pCAF co-cultures (Figure 6A&B). The most pronounced effect was observed with the FOLFIRINOX–trastuzumab combination. To exclude the possibility that these effects were driven by baseline differences in cancer cell numbers between WT and NFATC2+ pCAF conditions, responses were normalized to vehicle-treated controls. This analysis confirmed a synergistic interaction between FOLFIRINOX and trastuzumab specifically in NFATC2+ co-cultures (Figure 6C). These findings were further validated by flow cytometry–based quantification of cancer cell populations following combination treatments (Figure 6D).

**Figure 6.**
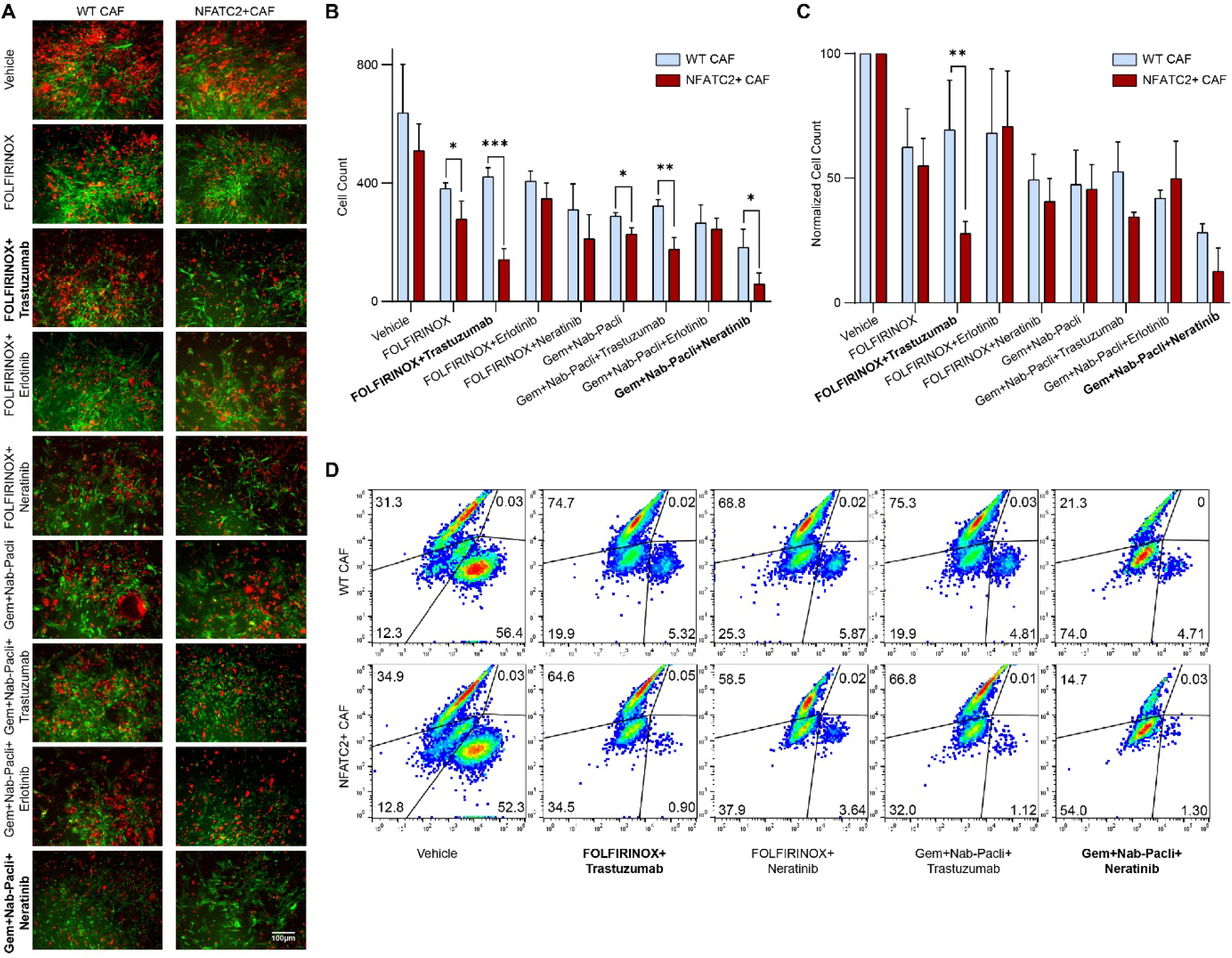
ERBB-targeted therapies enhance chemotherapy efficacy in NFATC2+ pCAF co-culture system. (A) Representative fluorescence microscopy images of PANC1-pCAF co-culture system under different treatment conditions. Cancer cells (mCherry+, red) and CAFs (eGFP+, green) were treated with vehicle control, standard chemotherapy regimens (FOLFIRINOX or Gemcitabine plus Paclitaxel) alone, or chemotherapy combined with ERBB-targeted inhibitors including Trastuzumab, Erlotinib and Neratinib. Scale bar, 10μm. (B) Quantification of PANC1 cancer cell numbers from fluorescence microscopy analysis. Data represent mean ± SEM from three independent experiments. Statistical significance: *p < 0.05, **p < 0.01, ***p < 0.001, t-test. (C) Normalized cell numbers of PANC1 from fluorescence microscopy analysis based on vehicle control from each group. Data represent mean ± SEM from three independent experiments. Same statistical significance marks used as (B). (D) Flow cytometry analysis of the co-culture system after three-day treatment. Density plots display mCherry fluorescence (x-axis) versus eGFP fluorescence (y-axis), revealing distinct cell populations: PANC1 cells (mCherry+, eGFP-, lower right quadrant), CAFs (mCherry-, eGFP+, upper left quadrant), and cellular debris (lower left quadrant). Quantification of PANC1 cell proportions demonstrating a marked reduction in the cancer cell population following chemotherapy combined with ERBB-targeted inhibitors, particularly Trastuzumab and Neratinib, in the presence of NFATC2+ pCAFs.

Collectively, these results demonstrate that the tumor-suppressive effects associated with NFATC2+ CAF states can be exploited in preclinical models through rational combination strategies targeting the ERBB signaling pathway. Overall, these findings suggest that NFATC2 expression in the tumor stroma may be used for stratification of tumors that are more likely to benefit from ERBB-targeted therapies in combination with standard chemotherapy.

### Validation of NFATC2 effect on chemotherapy response in another PDAC cohort as well as in rectal cancer

To validate and extend the link between NFATC2 expression and chemotherapy response, we analyzed an independent scRNA-seq dataset from PDAC patients treated with neoadjuvant chemotherapy as an external validation cohort^24^. UMAP analysis of CAF populations revealed distinct transcriptional states associated with treatment status (Figure 7A) and treatment response (Figure 7B). In line with our findings from the primary cohort, NFATC2 expression in CAFs was markedly higher in chemotherapy-treated patients compared to treatment-naïve tumors (Figure 7C). Moreover, NFATC2 expression was enriched in partial responders following neoadjuvant therapy compared untreated or progressive disease (Figure 7D), supporting treatment-associated activation of NFATC2-linked stromal programs.

**Figure 7.**
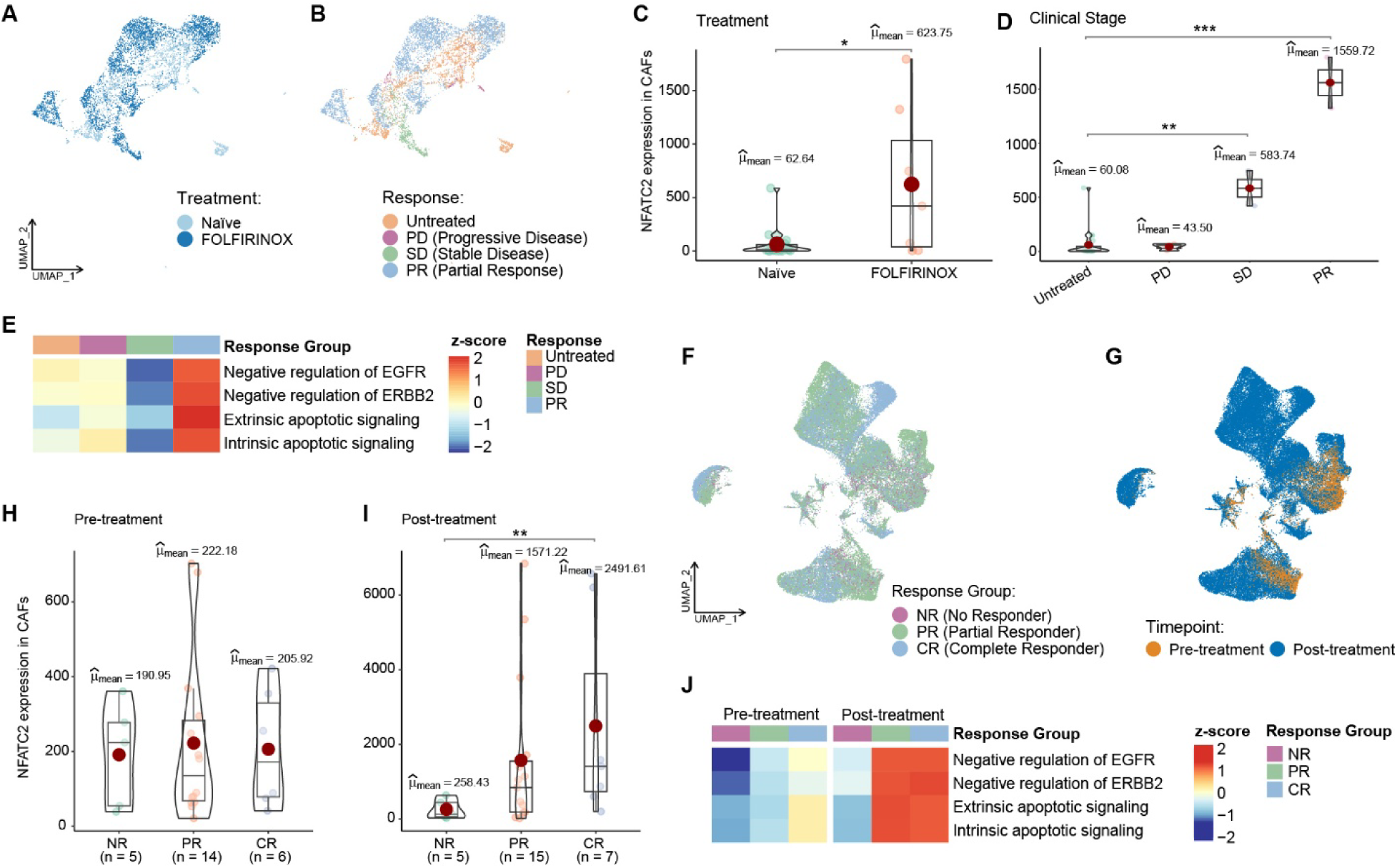
External data validation cohorts demonstrate that NFATC2 expression in CAFs is associated with chemotherapy response and treatment sensitivity across cancer types. (A) UMAP visualization of CAFs from an independent pancreatic cancer validation cohort^24^, colored by treatment status: untreated (light blue) and treated (dark blue) patients showing distinct transcriptional states between chemotherapy-exposed and treatment-naïve stromal populations. (B) UMAP visualization of the same CAFs colored by treatment response categories: untreated (orange), progressive disease (purple), stable disease (green), and partial response (blue), demonstrating response-associated distribution patterns of CAF subpopulations. (C) NFATC2 expression levels in CAFs stratified by treatment status, showing markedly elevated expression in chemotherapy-treated patients (μmean = 623.75, n=7) compared to untreated patients (μmean = 62.64, n=20) (p<0.05, t-test). (D) NFATC2 expression in CAFs stratified by treatment response, showing higher NFATC2 expression in partial response compared untreated or progressive disease (* p<0.05, ** p<0.01, *** p<0.001, t-test). (E) GSVA heatmap showing pathway enrichment scores in cancer cells from the same pancreatic cancer validation cohort^24^, stratified by treatment response. Rows represent different Gene Ontology Biological Process (GOBP) pathways including negative regulation of EGFR, ERBB signaling pathway, extrinsic apoptotic signaling, and intrinsic apoptotic signaling. Columns represent different treatment responses (Untreated, PD, SD, PR) with color intensity indicating pathway enrichment scores. (F) UMAP visualization of CAFs from a rectal cancer cohort^57^ colored by treatment response: no responder (NR, purple), partial responder (PR, green), and complete responder (CR, blue), showing distinct clustering patterns associated with therapeutic outcomes. (G) UMAP visualization of the same CAFs colored by timepoint: pre-treatment (orange) and post-treatment (blue), illustrating treatment-associated shifts in CAF transcriptional states. (H) NFATC2 expression levels in CAFs from the rectal cancer cohort prior to treatment, stratified by response group. (I) Violin plot displaying post-treatment NFATC2 expression levels in CAFs across treatment response groups, showing higher NFATC2 expression in complete response group (** p<0.01, t-test). (J) GSVA heatmap showing pathway enrichment scores in cancer cells from the rectal cancer cohort^57^, stratified by treatment response groups (NR, PR, CR) and timepoint (pre- and post-treatment). The same GOBP pathways are analyzed as in panel E, showing dynamic changes in pathway activities across response categories and treatment timepoints.

To determine whether CAF-associated NFATC2 enrichment was accompanied by transcriptional changes in malignant epithelial cells, we next examined pathway activity in cancer cells from the same cohort. Pathway-level GSVA revealed stage-dependent transcriptional programs, with apoptotic signaling preferentially enriched in advanced stages that typically receive more intensive chemotherapy (Figure 7E). Notably, negative regulation of EGFR and ERBB2 signaling pathways was also observed in chemotherapy-treated stages, particularly in best-responding group (partial response cases), consistent with our primary findings linking NFATC2+ pCAFs to suppression of ERBB signaling. Together, these data support an association between NFATC2 expression in CAFs, attenuation of ERBB-related programs in cancer cells, and enhanced treatment response.

To assess whether this association extends beyond pancreatic cancer, we analyzed a rectal cancer cohort^57^ treated with neoadjuvant FOLFIRINOX and stratified by treatment response. CAF clustering revealed distinct transcriptional patterns associated with treatment status and response (Figure 7F&G). Prior to therapy, NFATC2 expression levels were comparable across different response groups (Figure 7H), indicating that baseline NFATC2 expression does not predict therapeutic response. In contrast, post-treatment analysis revealed a strong association between NFATC2 expression and treatment efficacy, with the highest NFATC2-levels observed in complete responders and the lowest levels in non-responders (Figure 7I). Consistent with the PDAC cohorts, GSVA analysis further demonstrated dynamic regulation of apoptotic and ERBB-related pathways associated with treatment response and timepoint in cancer cells (Figure 7J), with complete responders showing enhanced apoptotic signaling and negative regulation of ERBB pathways following treatment.

To investigate the broader clinical relevance of NFATC2 beyond pancreatic cancer, we performed survival analysis across different tumor types using TCGA datasets. Consistent with our observations in PDAC (c.f. Supplementary Figure 2D), elevated NFATC2 expression was associated with improved overall survival in several malignancies, including lung adenocarcinoma, breast cancer, and head-neck squamous carcinoma (Supplementary Figure 6). These associations are concordant with our functional data and support a tumor-suppressive role for NFATC2 in specific TME contexts.

Collectively, these independent validation analyses demonstrate that NFATC2 expression in CAFs is induced by chemotherapy and consistently associates with favorable treatment response and conserved transcriptional programs across two different cancer types treated with similar neoadjuvant regimens. Together with our functional studies, these findings support NFATC2 as a stromal biomarker associated with chemotherapy sensitivity and suggest that tumors enriched for NFATC2+ CAFs may represent contexts in which ERBB-targeted therapies could be rationally combined with standard chemotherapy to improve therapeutic efficacy while potentially reducing treatment-associated toxicity.

## DISCUSSION

Cancers are organ-specific and driven by distinct mutational landscapes^58,59^, prompting extensive efforts to develop tumor type-specific targeted therapies. In contrast, cells of the TME are largely non-malignant and may rely on regulatory mechanisms shared across cancer types, potentially creating opportunities for more generalizable therapeutic strategies. Recent single-cell studies have shown that CAFs across multiple tumor types adopt recurrent functional states, including inflammatory and myofibroblastic phenotypes^23,60,61^. Understanding these CAF regulatory states is particularly important in PDAC due to its dense desmoplastic stroma with pronounced CAF heterogeneity and context-dependent effects on tumor progression and therapy response^5,10,62^. Attempts to therapeutically target CAFs have produced mixed outcomes^16^, with either tumor restraining or promoting effects depending on context and cell state^61^. For example, depletion of FAP⁺ CAFs has been associated with improved survival, whereas depletion of αSMA⁺ CAFs resulted in accelerated disease progression^21^, underscoring the need to distinguish tumor-promoting from tumor-restraining stromal states.

Building on this framework, we identified NFATC2+ CAFs as a treatment-responsive fibroblast subpopulation with tumor-suppressive properties in PDAC. Through integrated single-cell analysis, GRN modeling, and functional validation, we demonstrate that NFATC2+ CAFs associate with enhanced chemotherapy sensitivity, ERBB pathway suppression, activation of pro-apoptotic programs, and improved patient survival.

Transcriptional adaptation is a central feature of chemotherapy response, and our data highlight CAF regulatory states as a critical component of this process. GRN analysis revealed NFATC2 as a key TF selectively activated in CAFs of neoadjuvantly treated tumors, in contrast to the pro-fibrotic factors predominant in untreated tumors. This pattern suggests that therapeutic pressure drives reprogramming of stromal regulatory landscapes, in agreement with recent reports linking stromal features in PDAC to treatment responses ^21,28–30,62,63^. Although NFATC2+ CAFs share features with iCAFs, their strong association with treatment response, ERBB suppression, and pro-apoptotic signaling supports a functionally distinct, therapy-conditioned stromal state that emerges in response to chemotherapy, rather than a pre-existing baseline CAF subtype. This is consistent with recent perturbation-based studies showing that CAF phenotypes can be actively redirected from TGF-β–driven myofibroblastic programs toward interferon-responsive, anti-tumoral states with enhanced immune engagement^64^.

The association of NFATC2+ CAFs with better treatment response is further reflected in their enrichment for reactive subTME programs previously linked to immune infiltration and treatment sensitivity^20^. Consistently, downregulation of CXCL12 observed in NFATC2+ CAFs has been implicated in improved responsiveness to immunotherapy^49^ and suppression of myCAF states linked to enhanced T cell infiltration^64^. Notably, NFAT signaling has also been linked to inflammatory CAF programs driven by IL-1 pathway activation in PDAC^65^, whereas our results highlight NFATC2 as a regulator of therapy-associated stromal reprogramming emerging upon neoadjuvant treatment. Together, these findings align with the established plasticity of CAF states and context-specific transcriptional adaptation in PDAC^45,52^. While the upstream signals inducing NFATC2 in CAFs remain to be fully defined, our data suggest that immune-derived signals in treated tumors, particularly from T cells, may contribute to the emergence of NFATC2+ CAF states. This is supported by a previous report demonstrating a T cell-derived chemokine switch that promotes immune-activating CAF programs^66^, linking therapy-associated immune activity to stromal transcriptional reprogramming.

Experimental activation of NFATC2 in CAFs enhanced the sensitivity of cancer cells to FOLFIRINOX and other chemotherapeutic regimens, along with increased apoptotic activity. These effects were linked to coordinated modulation of downstream signaling pathways, such as suppression of ERBB signaling, a known contributor to PDAC progression and treatment resistance^67^, as well as promotion of epithelial containment and matrix anchorage of cancer cells, features associated with reduced invasive potential. Together, these findings position NFATC2 not merely as a marker of CAF heterogeneity, but as a transcriptional hub capable of rewiring stromal regulatory networks in response to therapeutic pressure.

Building on these NFATC2-associated changes in ERBB signaling and apoptotic priming, we reasoned that ERBB-targeted therapies might be particularly effective in NFATC2-enriched stromal contexts. Our combinatorial drug testing revealed distinct therapeutic vulnerabilities associated with NFATC2+ pCAF-mediated stromal reprogramming. In NFATC2+ pCAF co-cultures, HER2-targeting agent trastuzumab showed greater efficacy than the EGFR inhibitor erlotinib, suggesting preferential modulation of HER2-dependent signaling rather than uniform effects across the ERBB family. This is notable given the limited clinical success of EGFR/ERBB inhibitors in PDAC when applied in unselected patient populations^53,55,67^, highlighting a context-dependent sensitivity that becomes evident only when stromal states are taken into account. This preferential sensitivity may reflect several, non-mutually exclusive mechanisms, including selective interference with HER2/HER3 signaling, and lowering of the apoptotic threshold under chemotherapy-induced stress. Together, these findings suggest that NFATC2+ pCAF–associated stromal states can shape selective therapeutic sensitivities and support stratified combination strategies in PDAC.

Elevated NFATC2 expression was consistently associated with improved clinical outcomes across multiple cancer types with prominent EGFR/ERBB signaling and stromal involvement. Together with the treatment-associated NFATC2 upregulation and enhanced chemotherapy sensitivity observed in several PDAC cohorts as well as in rectal cancer, these findings support NFATC2 as a treatment-associated stromal biomarker with clinical relevance beyond pancreatic cancer. The convergence of prognostic relevance and treatment-induced NFATC2 activation suggests that NFATC2+ pCAF-mediated programs may represent a common adaptive stromal response to effective cytotoxic therapy. Accordingly, NFATC2-guided stratification could identify tumors more likely to benefit from chemotherapy and ERBB-targeted combinations, enabling the rational optimization of combination treatment strategies across selected solid tumor types.

Several considerations will be important for translating these findings toward clinical applications. While our in vitro co-culture experiments provide strong functional evidence for the tumor-suppressive role of NFATC2+ pCAFs, in vivo validation using complementary model systems will be required to establish how these stromal states function within complex tumor ecosystems. Further elucidation of the upstream cues that induce NFATC2 expression in CAFs during therapy, particularly immune-mediated signals, represents an important direction for future investigation. Moreover, the pronounced spatial heterogeneity of PDAC highlights the importance of spatially resolved analyses to determine how NFATC2+ CAFs are distributed across tumor regions, disease stages, and treatment contexts. Whether NFATC2 can be detected via minimally invasive approaches such as plasma-based assays remains to be explored but would substantially enhance biomarker feasibility. The long-term stability and persistence of NFATC2+ pCAFs following treatment cessation also remain unknown warranting longitudinal studies, which are challenging in pancreatic cancer given the poor patient survival. Despite these challenges, the combination of ERBB-targeted therapies with chemotherapy in NFATC2+ stromal contexts is particularly compelling, as it leverages clinically approved agents and could thus accelerate translational implementation.

In conclusion, identification of NFATC2+ pCAFs elucidates stromal heterogeneity, transcriptional adaptation, and TF-driven regulatory states in PDAC. Rather than universally targeting CAFs, our findings identify a treatment-responsive CAF subpopulation with tumor-suppressive properties, underscoring the need for precision therapeutic strategies that preserve beneficial stromal cells while avoiding non-selective CAF targeting. By defining a TF-driven stromal program linked to therapeutic benefit, this study provides mechanistic insight into therapy-induced stromal reprogramming and establishes a rational framework for precision medicine strategies that integrate stromal state into treatment stratification.

## Supporting information

Figure abstract

## RESOURCE AVAILABILITY

### Lead contact

Further information and requests for resources and reagents should be directed to and will be fulfilled by the lead contact, Biswajyoti Sahu.

### Materials availability

This study did not generate new unique reagents.

### Data and code availability

The primary single-nucleus RNA sequencing dataset analyzed in this study was previously published and is publicly available through the Gene Expression Omnibus (GEO) under accession number GSE GSE202051^34^. Validation cohort single-cell RNA sequencing data for PDAC is available under GEO accession GSE205013^24^, and rectal cancer scRNA-seq data is available from China National GeneBank Database (CNGBdb) with accession number CNP0004138^57^. Bulk RNA-seq data from pCAF-PANC1 co-culture experiments and conditioned medium treatments generated in this study have been deposited in the Gene Expression Omnibus (GEO) under accession number GSE316877 and are publicly available as of the date of publication. This study did not generate unique computational code. All analysis were performed using publicly available software and packages.

## ACKNOWLEDGMENTS

We thank Norwegian Sequencing Centre, Oslo, Norway for NGS services and the computational infrastructure at NCMBM for data analysis. B.S. was supported by Research Council of Norway (187615), Helse Sør-Øst, University of Oslo through the Norwegian Centre for Molecular Biosciences and Medicine (NCMBM) (to Sahu group), Norwegian Cancer Society (274630), South-Eastern Norway Health Authority (HSØ) (2025083), Sigrid Jusélius Foundation, Finnish Cancer Foundation and Jane and Aatos Erkko Foundation. T.A. was supported by the Research Council of Finland (grants 344698, 345803, and 367855); the Cancer Foundation of Finland, the Norwegian Cancer Society (grants 216104 and 273810), South-Eastern Norway Regional Health Authority (grants 2020026 and 2023105), Radium Hospital Foundation, the Sigrid Jusélius Foundation, and iCAN – Digital Precision Cancer Medicine Flagship (iCAN-MULTIDRUG). We thank Dr. Divyesh Patel with help in laboratory work and Dalila Sabrine Hedhili, Nann Kristoffersen for technical assistance. We thank Dr. Päivi Pihlajamaa and members of Sahu group for their inputs and scientific discussion.

## AUTHOR CONTRIBUTIONS

B.S. conceived and supervised the study. J.G. and S.C. performed in silico analyses. J.G. performed the experiments with support from C.W C.X and L.F. E.H. and S.D. contributed with the T cell isolation. C.V and T.A. contributed to data interpretation. B.S. and J.G. wrote the manuscript with contributions from all authors.

## DECLARATION OF INTERESTS

### STAR⍰METHODS

### KEY RESOURCES TABLE

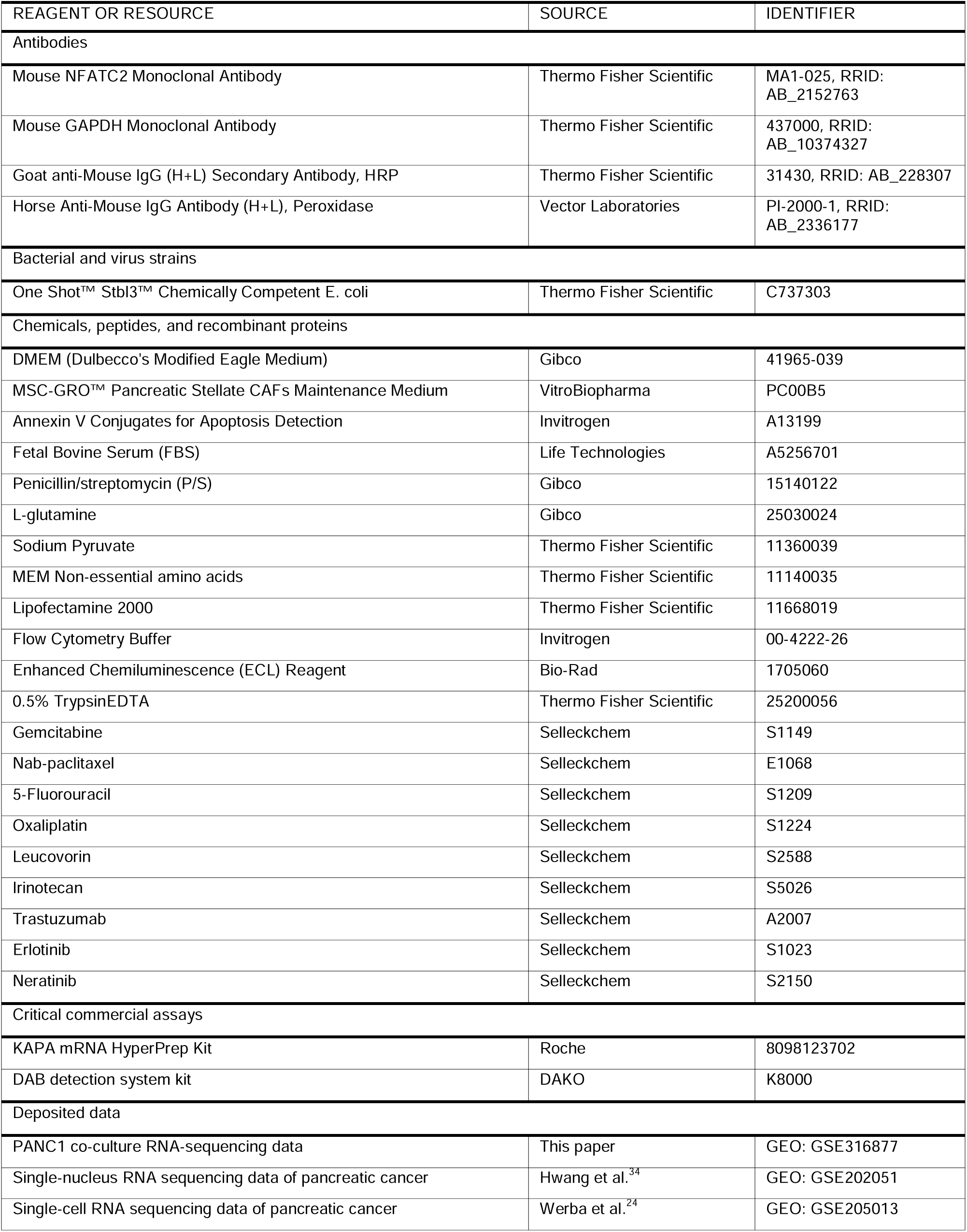
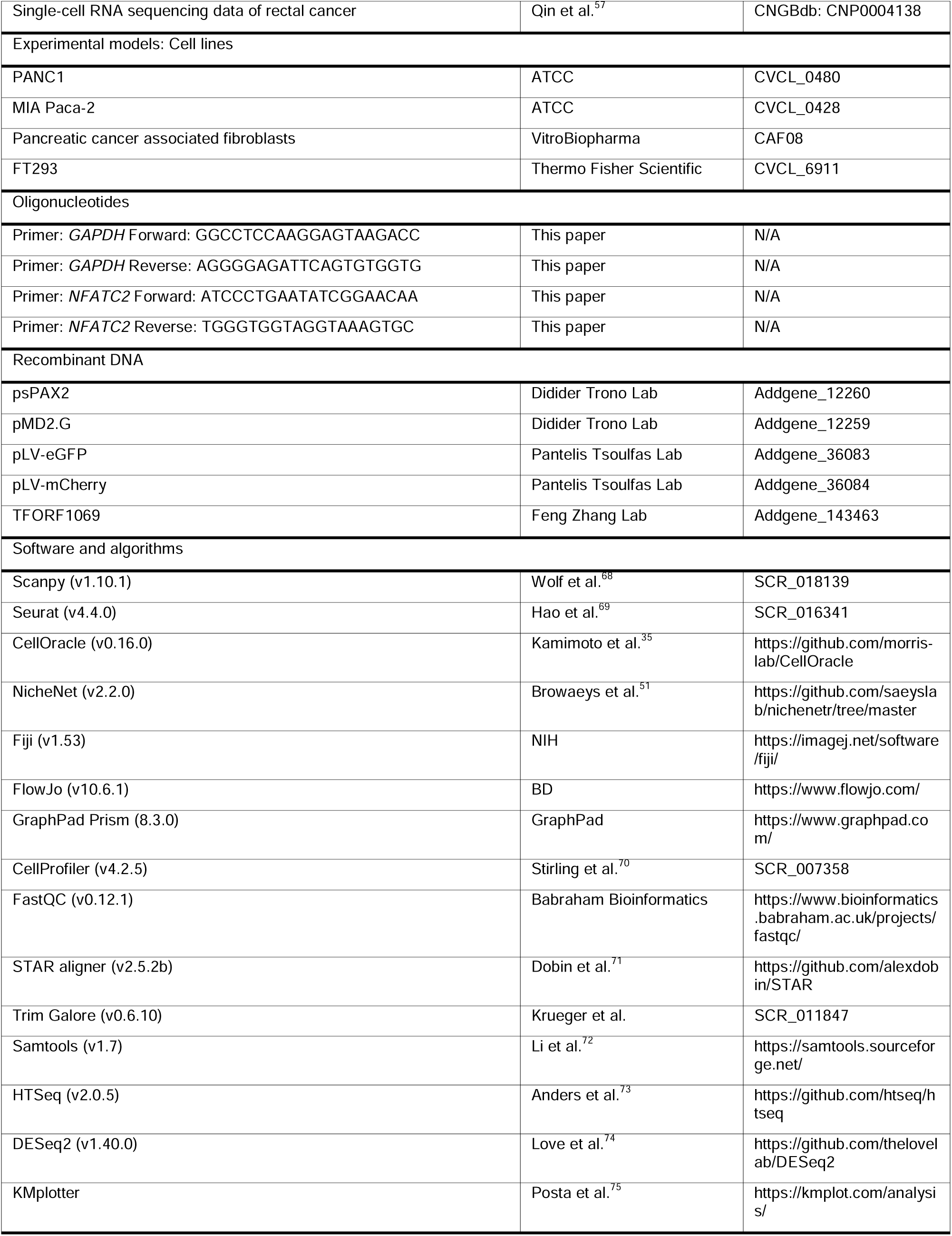

## METHOD DETAILS

### Single-nucleus sequencing data analysis

We analyzed a publicly available snRNA-seq dataset from pancreatic ductal adenocarcinoma patients^34^, comprising 42 samples (18 chemotherapy-treated and 24 treatment-naive). Raw count matrices were processed using Scanpy (v1.10.1, RRID: SCR_018139)^68^ in Python (3.9.22) and Seurat (v4.4.0, RRID: SCR_016341)^69^ in R (4.4.1). Nuclei were filtered based on the following criteria: 200-6000 detected genes, <20% mitochondrial gene content, and <1% hemoglobin gene content. Data were normalized using the standard log-normalization method, and the top 2000 highly variable genes were identified using the function ‘FindVariableFeatures’. Principal component analysis was performed on scaled data, and the top 30 principal components were used for downstream analysis. UMAP dimensionality reduction was applied for visualization. Leiden clustering (resolution = 0.5) was performed to identify cell clusters.

Major cell types were annotated as previously described in the original publication^34^, including epithelial cells, endothelial cells, immune cells (T cells, myeloid cells, B cells), and other stromal populations. Then, stromal cell populations (fibroblasts, pericytes, and vascular smooth muscle cells) were subset and re-clustered at higher resolution (resolution = 1.0) to identify distinct subtypes. CAF subtypes were classified based on established marker gene expression patterns^17,76,77^: Inflammatory CAFs (iCAFs): characterized by high expression of inflammatory markers including C3, C7, FBLN1 and IGF1. Myofibroblastic CAFs (myCAFs): defined by expression of FAP, ACTA2, INHBA and POSTN. Pericytes were characterized by RGS5, MCAM and TAGLN expression. Vascular smooth muscle cells (vSMCs) were defined by high expression of ACTA2, TAGLN and SMTN.

To validate our findings, we analyzed a second independent scRNA-seq dataset from PDAC patients^24^, comprising samples from both chemotherapy-treated (n=7) and treatment-naive (n=20) primary PDAC and liver metastasis tissues. This dataset was obtained through direct correspondence with the original authors. Raw count matrices were processed using Seurat (v4) in R (4.4.1) with identical quality control criteria as applied to the primary dataset: 200-6000 detected genes, <20% mitochondrial gene content, and <1% hemoglobin gene content. Data were normalized using standard log-normalization, and the top 2000 highly variable genes were identified. Cell type annotations provided by the original authors were utilized to examine gene expression patterns across treatment conditions and cellular populations.

For additional validation across cancer types, we analyzed a third independent scRNA-seq dataset from rectal cancer patients treated with neoadjuvant FOLFIRINOX^57^, containing 52 samples (25 treated and 27 treatment-Naïve) from 29 patients. The data was downloaded from China National GeneBank Database (CNGBdb) with accession number CNP0004138 and processed using the same computational pipeline and quality control thresholds.

### Construction of gene regulatory network (GRN) and identification of hub TFs in PDAC

To identify specific relations between TFs and genes, we created a gene regulatory network (GRN). This network allows to identify relations between TFs and target genes (TGs) at a deeper level, revealing dependencies between gene-gene regulation. The network is built using CellOracle (v0.16.0 on Python 3.8)^35^, a state-of-the-art tool specifically designed to generate GRNs for single cell datasets. As initial configuration, the tool requires a set of pre-existing TF-TG relations. This was created by using the well-known gimmemotifs (v5 vertebrate) human dataset^78^ provided by the authors of the tool. After this step, we split the single cell dataset into different subgroups of interest, like cell type and treatment information. After this initial sub-classification, the set of cells is reduced to allow a faster and more accurate computation of the network. We included 10k cells (randomly sampled) and 3k genes (highly variable genes) in each network. One network contained the cells coming from treated samples and the other contained cells coming from treatment-naive samples. Both networks used epithelial cells (malignant and benign) and CAF. Then the method uses Bayesian ridge regression model to predict the contribution of each TF to modulate a target gene expression. After this step, we have two distinct GRN that can be used to extract information about the connections of each TF and TG. By using common networks measures like the eigenvector centrality, we retrieved a set of TFs representing hubs (heavily connected genes). These hubs are heavily connected to genes with many connections in the network, meaning that the influence of these TF in the network is high.

### Differential gene expression analysis

Differential expression analysis was performed using a pseudobulk approach. First, malignant cells from each patient were aggregated by patient ID to generate pseudobulk expression profiles, which reduces single-cell noise while preserving biological variability between patients. Gene expression counts were summed across all malignant cells within each patient, and only genes expressed in at least 10% of patients were retained for downstream analysis. Pseudobulked samples were grouped according to neoadjuvant therapy response categories (untreated, poor response, minimal response, and moderate response) and analyzed using DESeq2^74^. Differentially expressed genes (DEGs) were identified based on following threshold: absolute log2 fold change ≥ 1 and adjusted p-value < 0.05.

### Functional pathway enrichment analysis

Gene Ontology biological process (GO BP) and Kyoto Encyclopedia of Genes and Genomes (KEGG) enrichment analyses were conducted using the R package ‘clusterProfiler’^79^. Threshold of p value < 0.05 was used for filtering. Gene set enrichment analysis (GSEA) and gene set variation analysis (GSVA) were implemented with the ‘GSVA’ package^80^, and visualized using the ‘GseaVis’ package. Gene sets used including Hallmark, GO and KEGG were obtained from MSigDB^81^.

### Survival analysis

Kaplan-Meier analysis was conducted using the R packages ‘survival’ and ‘survminer’. For single-cell data, patients were stratified into high and low NFATC2 expression groups based on the median pseudobulk expression level of NFATC2 in CAFs as the cutoff threshold. Kaplan-Meier survival curves were generated for high and low NFATC2 expression groups, and the log-rank test was used to assess statistical significance between survival curves, with p-values < 0.05 considered statistically significant.

Bulk RNA sequencing data from The Cancer Genome Atlas (TCGA) pancreatic adenocarcinoma cohort (PAAD, n=178) were obtained through UCSC Xena browser^82^. Patients were similarly stratified based on median NFATC2 expression levels in tumor samples for Kaplan-Meier analysis. The database ‘kmplotter’^75^ was utilized for following pan-cancer survival analysis.

### Cell crosstalk analysis

Ligand-receptor interaction analysis was performed using the R package ‘nichenetr’ ^51^. Cell types with sufficient cell numbers (>50 cells) were included in the analysis. Potential ligands were identified from sender cells (CAF subtypes) based on expression levels and cell type specificity. Ligand activity scores were calculated using NicheNet’s built-in ligand-target regulatory network.

### Immunohistochemistry

Human pancreatic cancer tissues were obtained from Oslo University Hospital patients undergoing neoadjuvant treatment (FOLFIRINOX, 4 cycles) followed by surgery (n=11) or upfront surgery (n=4). The collection of samples was approved by the Regional Ethics Committee (REK 2015/738). All patients had given written informed consent for the use of tumor tissue for research purposes.

Formalin-fixed paraffin-embedded 3.5-micron thick whole tissue sections were incubated with mouse anti-NFATC2 (1:25, overnight incubation, Thermo Fisher Scientific, #MA1-025, RRID: AB_2152763) following antigen-retrieval (95°C, 30 min, citrate buffer solution, pH 6.0). Subsequently, sections were incubated with horseradish peroxidase (HRP)-conjugated secondary antibody (Vector Laboratories, #31430, RRID: AB_2336177) for 30 minutes, stained with a DAB detection system kit (EnVision Flex DAB+; DAKO North America, Santa Clara, CA, USA), and counterstained with hematoxylin. Immunostaining was assessed on digital whole slide images by manual counting of positive and negative stromal cells in 5 hot spots (each measuring 0.6 mm^2^) containing at least 200 stromal cells. The percentage of positive stromal cells averaged over the 5 hot spots was compared between cases.

### Cell lines and plasmids

Human pancreatic carcinoma cell lines PANC1 (RRID: CVCL_0480), MIA PaCa-2 (RRID: CVCL_0428) were cultured in DMEM (Gibco, Life Technologies) supplemented with 2 mM L-glutamine (Gibco, 25030024), 10% fetal bovine serum (FBS), and 1% Penicillin-Streptomycin (Gibco, 15140122). Human pancreatic adenocarcinoma cancer-associated fibroblasts (VitroBiopharma, #CAF08) were cultured in MSC-GRO™ Pancreatic Cancer-Associated Fibroblast (CAF) Maintenance Medium (VitroBiopharma, #PC00B5) with 1% Penicillin-Streptomycin. 293FT (Thermo Fisher Scientific, #R70007, RRID: CVCL_6911) cells were cultured in DMEM supplemented with 10% FBS, 6mM L-glutamine (Gibco, 25030024), 1% Penicillin-Streptomycin, 1% Sodium Pyruvate (Thermo Fisher Scientific, #11360039) and 1% MEM Non-essential amino acids (Thermo Fisher Scientific, #11140035). All cell lines were tested for mycoplasma contamination, cultured at low passages and maintained in a humidified incubator at 37 °C and with 5% CO2. NFATC2 full-length ORF plasmid was a gift from Feng Zhang (Addgene, #143463, RRID: Addgene_143463)^83^.

### Lentivirus production and stable transduction

For virus production, NFATC2 full-length ORF plasmid was co-transfected with the packaging plasmids psPAX2 (Addgene #12260, RRID: Addgene_12260) and pMD2.G (Addgene, #12259, RRID: Addgene_12259) in a ratio of 4:3:1 into 293FT cells with Lipofectamine 2000 (Thermo Fisher Scientific, #11668019). Then, culture medium was replenished on the following day and supernatant containing the viral particles was collected after 48 h ours, filtered with 0.45 mm filters (Merck, #SE1M003M00), and concentrated using Lenti-X concentrator (Clontech, #631232), tittered using qPCR Lentivirus Titer Kit (ABM, # LV900) following the manufacturer’s protocol and stored as single-use aliquots in-80 °C.

For stable transduction, an empty vector was transfected as control. A total of 4 × 10^5^ pCAFs in 2 mL of medium with 2 µg/mL polybrene were infected with 30 uL of lentivirus supernatant. After 72 h, 2 μg/ml puromycin was used for NFATC2 overexpression selection of a stable cell pool. Cell sorting was used for eGFP and mCherry fluorescent marker selection. The efficiency of overexpression in a stable cell line was validated by qRT-PCR and Western blot assays as described in following methods.

### Western blot analysis

Pancreatic CAFs were lysed in RIPA Buffer (Thermo Fisher Scientific, #89900) supplemented with protease inhibitor cocktail (Thermo Fisher Scientific, #78441). Protein concentration was determined using the Bradford Protein Assay Kit (Bio-Rad, #5000202). Equal amounts of protein (20 μg) were separated by SDS-PAGE and transferred onto PVDF membranes (Bio-Rad, #1620177). Membranes were blocked with 5% non-fat milk in TBST (Tris-buffered saline with 0.1% Tween-20) The membranes were then incubated overnight at 4°C with primary antibodies diluted in 1% BSA in TBST: anti-NFATC2 (1:1000, Thermo Fisher Scientific, #MA1-025, RRID: AB_2152763) and anti-GAPDH (1:5000, Thermo Fisher Scientific, #437000, RRID: AB_10374327) as a loading control. Following three washes with TBST, membranes were incubated with Goat anti-Mouse Secondary antibody (1:10000, Thermo Fisher Scientific, #31430, RRID: AB_228307) for 1 hour at room temperature. Protein bands were visualized using enhanced chemiluminescence (ECL) reagent (Bio-Rad, #1705060) and detected with a ChemiDoc imaging system (Bio-Rad, #12003153).

### Direct co-culture of cancer cells and pCAFs and drug treatment

Prior to establishing the co-culture system, we performed ratio optimization experiments to determine the optimal CAF: cancer cell ratio according to previous reports^45–47^. PANC1 cells were co-cultured with pancreatic CAFs at four different ratios: 3:1, 2:1, 1:1, and 1:3 (CAF: cancer cell). After 72 hours of co-culture, we found that the 3:1 ratio maintained the most stable co-culture conditions without overgrowth of either cell type. Therefore, a ratio of 3:1 was selected for all subsequent co-culture experiments. Drug treatment was initiated on the second day following cell seeding, and subsequent quantitative and functional assays were performed on the fourth day.

Drug treatments were administered at the following concentrations^84–87^. All drugs were dissolved in DMSO and diluted to ensure a final DMSO concentration of ≤1% in each well. FOLFIRINOX: 5-Fluorouracil (Selleckchem, #S1209) at 10 μM, oxaliplatin (Selleckchem, #S1224) at 100 nM, leucovorin (Selleckchem, #S2588) at 132 nM and irinotecan (Selleckchem, #S5026) at 123 nM. Other combination therapy: Gemcitabine (Selleckchem, #S1149) at 100 nM, nab-paclitaxel (Selleckchem, #E1068) at 10 nM, trastuzumab (Selleckchem, #A2007) at 10 μg/mL, erlotinib (Selleckchem, #S1023) at 1 μM, and neratinib (Selleckchem, #S2150) at 100 nM.

### Image acquisition and quantitative analysis

Fluorescence images were acquired using Zeiss AxioVert fluorescence microscope. For each experimental condition, at least 3-4 random fields of view were captured per well, multiple with 3 biological replicates. Images were acquired with consistent exposure times and laser power settings across all samples to ensure comparability.

Fluorescence microscopy images from co-culture experiments were quantitatively analyzed using CellProfiler^70^ (version 4.2.5, RRID: SCR_007358). Images were acquired in CZI format containing dual fluorescence channels: mCherry-labeled cancer cells (red) and eGFP-labeled pCAFs (green). Raw images were imported into CellProfiler and underwent the following preprocessing steps and parameters: Multi-channel fluorescence images were separated into individual channels using the “ColorToGray”. Then, “CorrectIlluminationApply” module was used to correct for uneven background illumination. Noise reduction with Gaussian filtering (sigma = 2-3 pixels). Total cell numbers for each fluorescence channel (mCherry+ cancer cells and eGFP+ CAFs) were quantified per field of view.

### RNA isolation, quantitative real-time PCR and RNA-sequencing

Total RNA was isolated from two experimental designs: (1) direct co-culture of PANC1 with WT CAF and NFATC2+ CAF for 3 days with or without FOLFIRINOX treatment, (2) PANC1 cultured with both CAF-conditioned medium for 3 days with or without FOLFIRINOX treatment. RNA extraction was performed using the RNeasy Mini Kit (Qiagen, #74136) according to the manufacturer’s protocol. RNA concentration and purity were assessed using a NanoDrop spectrophotometer, with A260/A280 ratios between 1.8-2.0 considered acceptable. NFATC2 expression was quantified using the following primer sequences (forward: 5’-ATCCCTGAATATCGGAACAA-3’, reverse: 5’-TGGGTGGTAGGTAAAGTGC-3’). GAPDH served as the internal reference gene (forward: 5’-GGCCTCCAAGGAGTAAGACC-3’, reverse: 5’-AGGGGAGATTCAGTGTGGTG-3’). Relative gene expression was calculated using the 2(-ΔΔCt) method, with each sample analysed in triplicate.

RNA-seq libraries were constructed using 800-1000 ng of total RNA with the KAPA mRNA HyperPrep Kit (Roche, #8098123702) following the manufacturer’s protocol. Libraries were quantified using Qubit dsDNA HS Assay Kit. Libraries were sequenced on Illumina NovaSeq X platform generating 150 bp paired-end reads.

Raw sequencing reads were assessed for quality using FastQC (v0.12.1). Adapter sequences and low-quality bases were trimmed using Trim Galore (v0.6.10) with a quality threshold of Phred score ≥20 and minimum read length of 20 bp. Quality-filtered reads were aligned to the human reference genome (GRCh38) using STAR aligner (v2.5.2b)^71^ in two-pass mode with default parameters. Uniquely mapped reads were retained for downstream analysis. The resulting BAM files were sorted and indexed using Samtools (v1.7)^72^. Gene-level read counts were quantified using HTSeq (v2.0.5)^73^ in union mode with GENCODE v38 gene annotations. Differential gene expression analysis was performed using DESeq2 (v1.40.0)^74^ in R (v4.3.0). Raw count matrices were normalized using the DESeq2 median-of-ratios method to account for library size and composition biases. Genes with fewer than 10 reads across all samples were excluded from analysis. Statistical significance was determined using the Wald test with Benjamini-Hochberg correction for multiple testing.

### Flow cytometry analysis

Following treatments for 3 days, co-cultured cells were harvested for flow cytometric analysis. Culture medium was aspirated from the co-culture systems, and cells were washed once with PBS. Cells were dissociated using 0.25% trypsin-EDTA solution and incubated at 37°C for 5 minutes. Trypsin activity was neutralized by adding an equal volume of complete medium containing FBS. Single-cell suspensions were generated by gentle pipetting, followed by centrifugation at 400g for 4 minutes at room temperature. Cell pellets were resuspended in 1 mL flow cytometry buffer (Invitrogen, #00-4222-26) and passed through 40 μm cell strainers (Corning, #CLS431750) to remove cell aggregates and debris. Flow cytometric analysis was performed using a Sony SH800S Cell Sorter (Sony Biotechnology) operating in analysis mode. The instrument was cleaned and calibrated according to manufacturer protocols before each experimental session. Standard fluorescent beads were used for instrument calibration and quality control. Single-stained controls (eGFP-positive CAFs only and mCherry-positive PANC1 cells only) and unstained cells were used as negative controls for compensation adjustment and gating strategy optimization. Cell suspensions were loaded into the flow cytometer sequentially, with data acquisition initiated immediately upon sample loading. Data acquisition was performed until the entire sample from every condition was processed to ensure comprehensive quantitative assessment of cell population ratios and amounts.

## SUPPLEMENTAL INFORMATION

**Document S1: Figures S1–S6.**

**Document S2: Table S1. NFATC2 immunohistochemistry quantification in PDAC tissue microarray. Related to Figure 2**.

**Table S2. Differential expressed gene list from stromal cell subtypes. Related to Figure 2**.

**Table S3-S6. Differential expressed genes of PANC1 in different co-culture conditions. Related to**

