## Supplementary figures for "NFATC2 in pancreatic cancer-associated fibroblasts predicts treatment response and facilitates ERBB-targeted therapies"

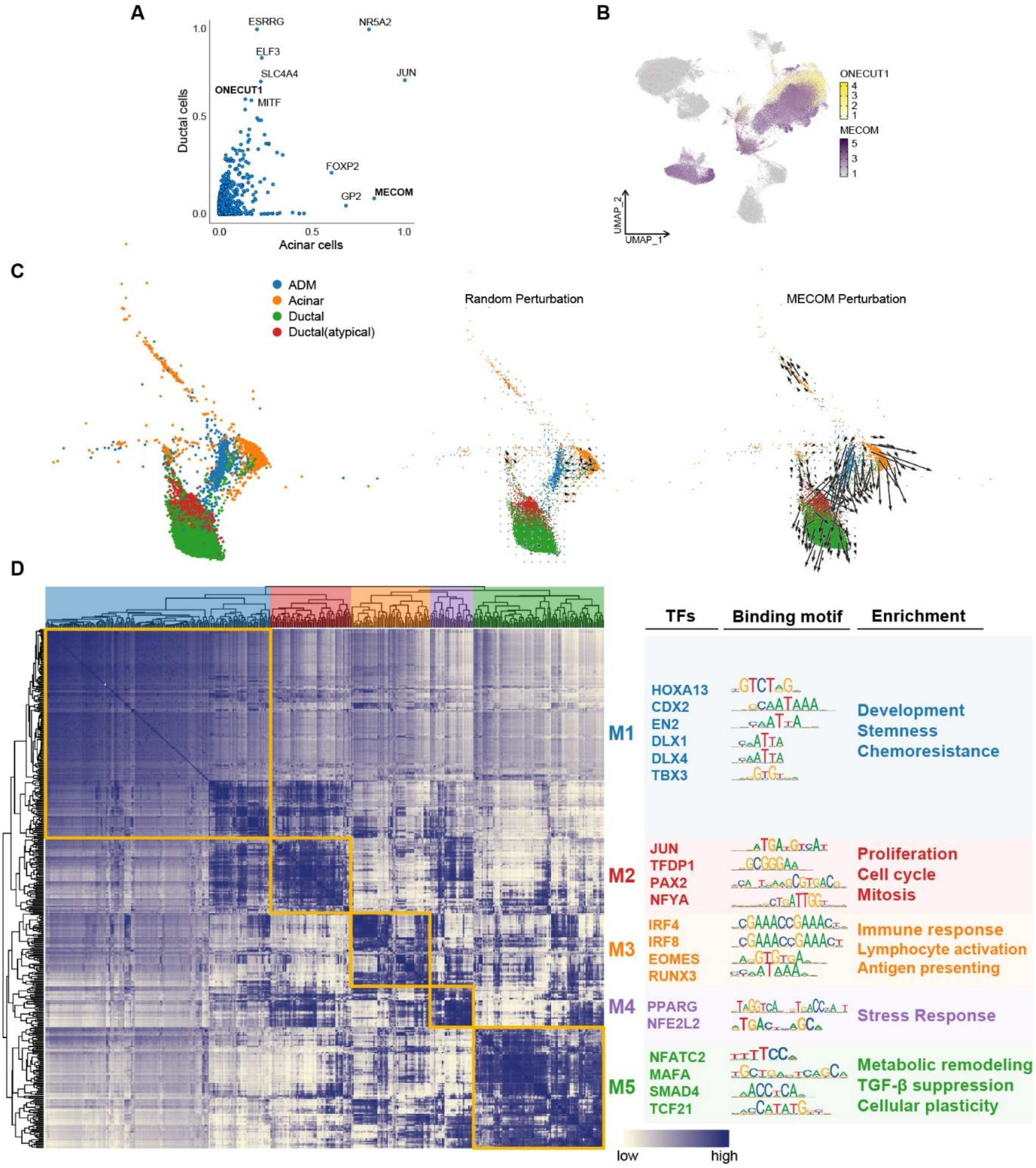


**Supplementary Figure 1. TF activity landscape across single-cell populations reveals functionally distinct regulatory modules. Related to Figure 1.**

(A) Comparison of eigenvector centrality scores between two pancreatic epithelial cell populations, acinar and ductal cells. This proof-of-concept analysis highlights the enrichment of MECOM in acinar cells and ONECUT1 in ductal cells.

(B) UMAP plot of single cells from 42 PDAC patients[1] colored by expression levels of ONECUT1 (top, yellow scale) and MECOM (bottom, purple scale), highlighting distribution patterns of key TFs across ductal and acinar cell populations.

(C) The effect of in-silico MECOM knockout on cell state transitions using CellOracle[2] Arrow plot shows that MECOM perturbation drives acinar cells away from the acinar-to-ductal metaplasia (ADM) cluster, whereas random perturbation as control has no effect on cell state transitions.

(D) Hierarchical clustering heatmap of TF activity across all cell types using SCENIC[3] identifies five distinct regulatory modules (M1-M5, outlined in yellow boxes). Columns represent individual cells grouped by cell type (colored bars at top), and rows represent TFs ordered by hierarchical clustering. Color intensity indicates activity level (blue = low, white = high). Each module is characterized by representative TFs, their consensus DNA binding motifs, and associated functional enrichments: M1 (blue) - development, stemness, and chemoresistance; M2 (red) - proliferation, cell cycle, and mitosis; M3 (orange) - immune response, lymphocyte activation, and antigen presentation; M4 (purple) - stress response; M5 (green) - metabolic remodeling, TGF-β suppression, and cellular plasticity. NFATC2 is enriched within M5, suggesting its role in metabolic and phenotypic reprogramming of CAFs.

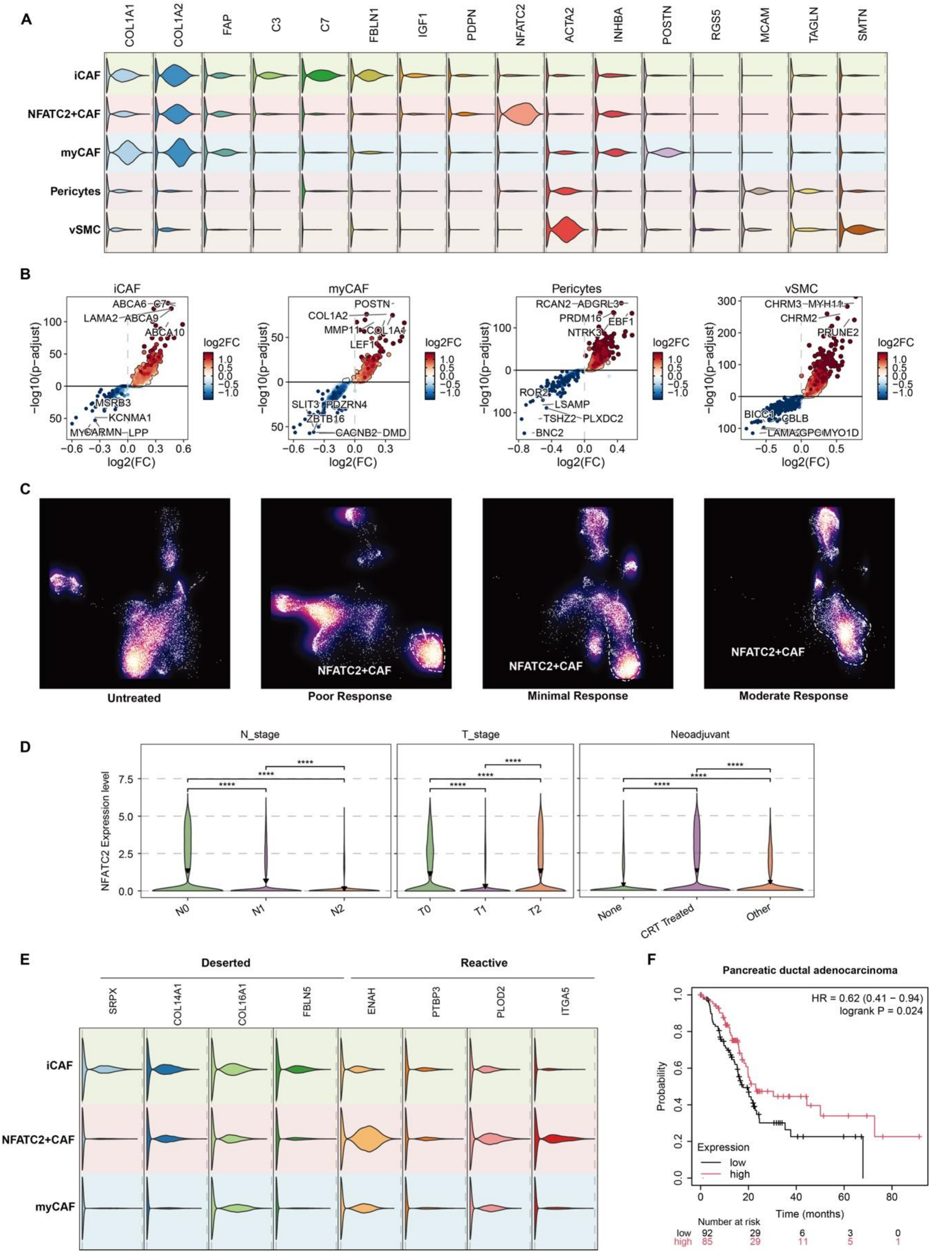


**Supplementary Figure 2. Differential gene expression and NFATC2 correlation with clinical parameters. Related to Figure 2.**

(A) Violin plots showing expression distribution of canonical stromal cell subtype markers across identified populations. Each plot displays the expression level of key markers for iCAF,

NFATC2+CAF, myCAF, pericytes, and vSMC (vascular smooth muscle cells) populations.

(B) Volcano plots showing differential gene expression in each stromal cell subtype compared to other subtypes (iCAF, myCAF, pericytes and sSMCs). Red dots: upregulated genes; blue dots: downregulated genes; gray: non-significant. Key marker genes are labeled.

(C) Density contour plots overlaid on UMAP showing spatial distribution patterns of cells from different response groups, with NFATC2+ CAF cluster (dotted outline) enriched in better responders.

(D) Violin plots showing NFATC2 expression distribution stratified by lymph node metastasis stage (N_stage: N0, N1, N2), tumor size stage (T_stage: T0, T1, T2), and neoadjuvant treatment status (None, CRT treated, Other). Each violin represents the probability density of NFATC2 expression levels, with wider sections indicating higher probability of samples at that expression value. Statistical significance was assessed using Kruskal-Wallis test, with pairwise comparisons indicated by asterisks (**** $P < 0.0001$).

(E) Violin plots showing expression of "deserted" versus "reactive" stroma markers (defined by ref. [20] as cited in the main text) across CAF subtypes. Left panel: Deserted markers (SPRX, COL14A1, COL16A1, FBLN1) showing higher expression in iCAFs. Right panel: Reactive markers (ENAH, FAP3, PLOD2, ITGA5) showing enrichment in NFATC2+CAFs. This demonstrates that NFATC2+CAFs exhibit a reactive stroma phenotype associated with enhanced tumor-stroma interactions.

(F) Kaplan-Meier survival curve showing overall survival stratified by NFATC2 expression levels (high vs low) from the TCGA pancreatic cancer dataset. It displays survival probability over time with hazard ratios (HR), 95% confidence intervals, and log-rank p-values. Red line represents high NFATC2 expression group, black line represents low NFATC2 expression group.

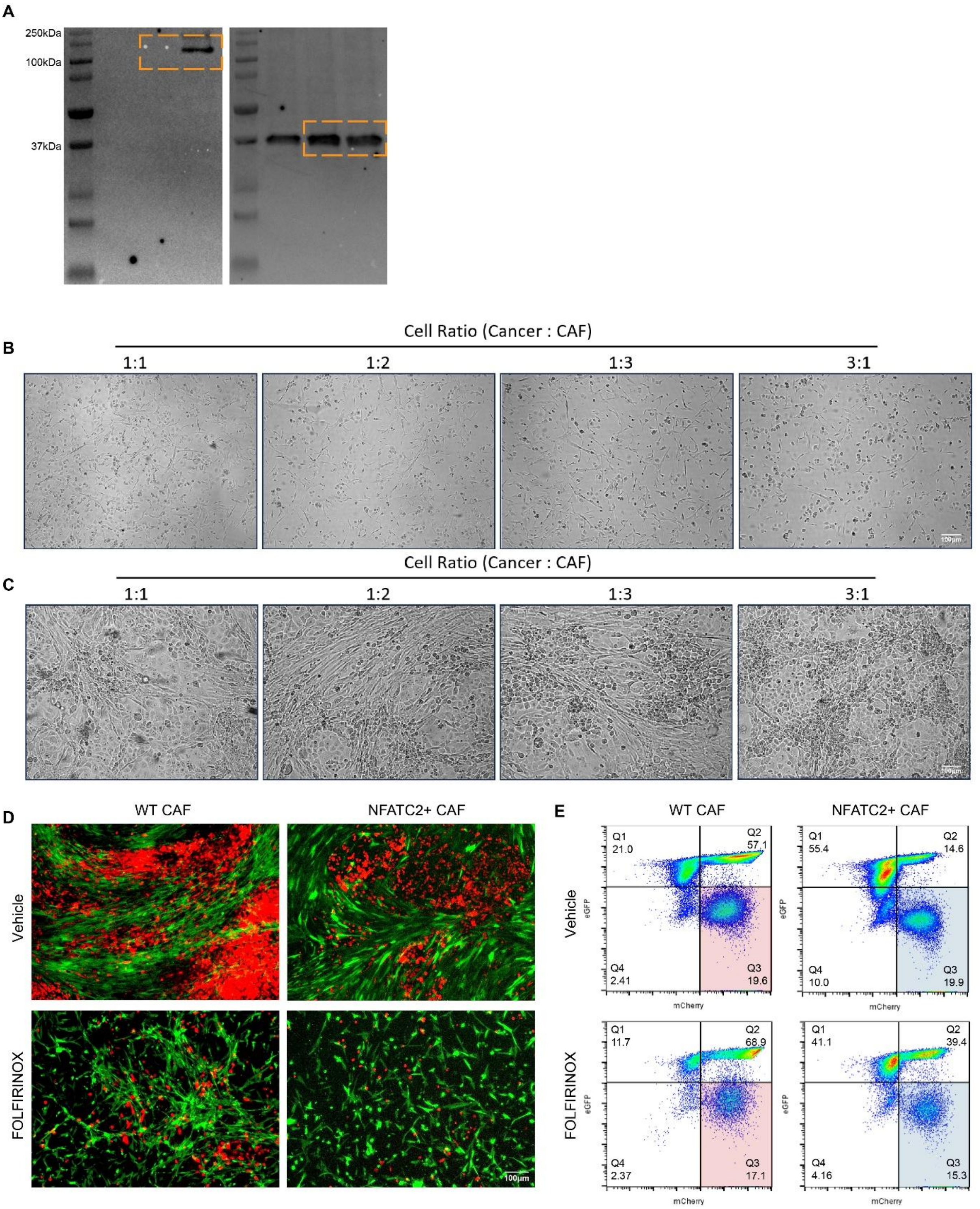


**Supplementary Figure 3. Co-culture optimization and validation of the results in MIA PaCa-2 cell line. Related to Figure 3.**

(A) Full-length Western blot image showing NFATC2 expression in WT pCAF (left and middle), and lentiviral NFATC2 overexpression (right). Orange boxes highlight NFATC2 band (~135 kDa, upper) and GAPDH loading control (~37 kDa, lower) presented in Figure 3A.

(B) Day 1 images of PANC1 cancer cells co-cultured with CAFs at ratios of 1:1, 1:2, 1:3, and 3:1 (CAF:cancer cell).

(C) Day 3 images of the same co-culture conditions, demonstrating progressive changes in cell density and morphology. The 3:1 ratio maintained optimal co-culture stability without overgrowth of either cell type.

(D) Representative fluorescence microscopy images of MIA PaCa-2 pancreatic cancer cell line (red, mCherry) co-cultured with wild-type CAFs (WT) or NFATC2-expressing (NFATC2+) CAFs (green, eGFP) under vehicle control (upper panels) or FOLFIRINOX treatment (lower panels). Scale bar, 100 µm.

(E) Flow cytometry analysis of MIA PaCa-2-CAF co-culture systems. Scatter plots displaying mCherry fluorescence (x-axis) versus eGFP fluorescence (y-axis) for the same experimental conditions as shown in panel C. Left panels show WT CAF and right panels NFATC2+ CAF co-cultures, with upper panels representing vehicle treatment and lower panels representing FOLFIRINOX treatment. Cancer cell populations (mCherry+eGFP-) are located in the lower right quadrant of each plot.

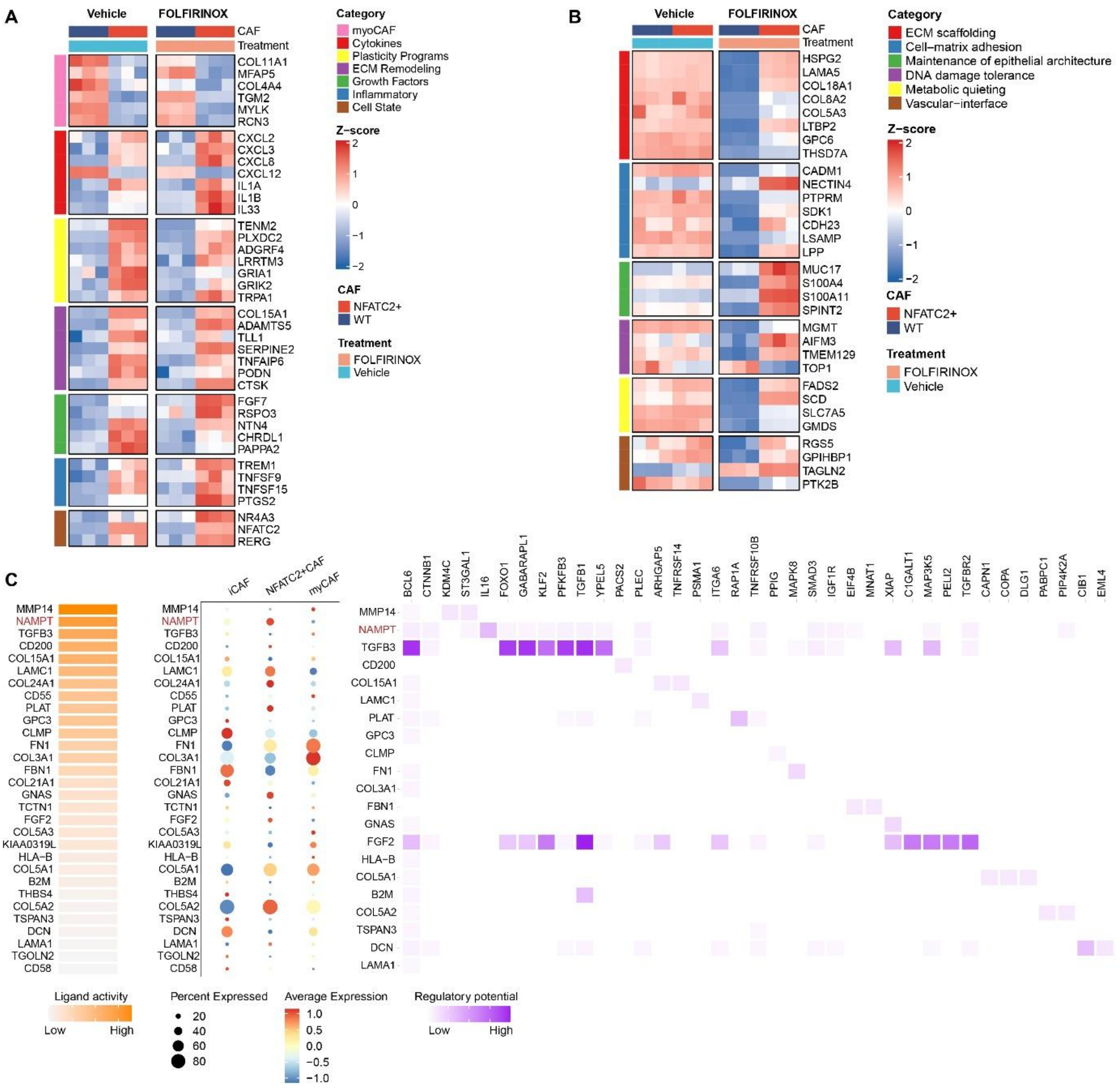


**Supplementary Figure 4. Transcriptome correlation and NicheNet-predicted ligand-receptor interactions between NFATC2+ CAFs and PANC-1 cells. Related to Figure 4.**

(A) Heatmaps showing Z-score normalized expression of selected genes in a direct co-culture system of PANC1 with either WT or NFATC2+ CAFs, treated with vehicle or FOLFIRINOX. Genes are grouped by functional categories: myCAF signature (pink), Cytokines (red), Plasticity Programs (yellow), ECM Remodeling (purple), Growth Factors (green), Inflammatory (blue), and Cell State (brown). Values for three replicate samples in each condition are shown with Z-score ranging from -2 (downregulated) to +2 (upregulated).

(B) Heatmaps showing Z-score normalized expression of genes in PANC-1 cells co-cultured with conditioned medium from either WT or NFATC2+ CAFs, treated with vehicle or FOLFIRINOX. Genes are grouped by functional categories: ECM Refolding (red), Cell-matrix adhesion (blue), Maintenance of epithelial architecture (green), DNA damage tolerance (purple), Metabolic quieting (yellow), and Vascular interface (brown). Layout and color scales are as in panel A.

(C) NicheNet ligand-receptor interaction analysis across CAF subtypes. Left panel shows ligand activity levels (orange gradient), percentage of cells expressing each ligand (dot size), and average expression levels (blue-to-red color scale) for key secreted across different CAF subtypes. Right panel displays predicted targets and the regulatory potential (purple gradient intensity) of these ligands.

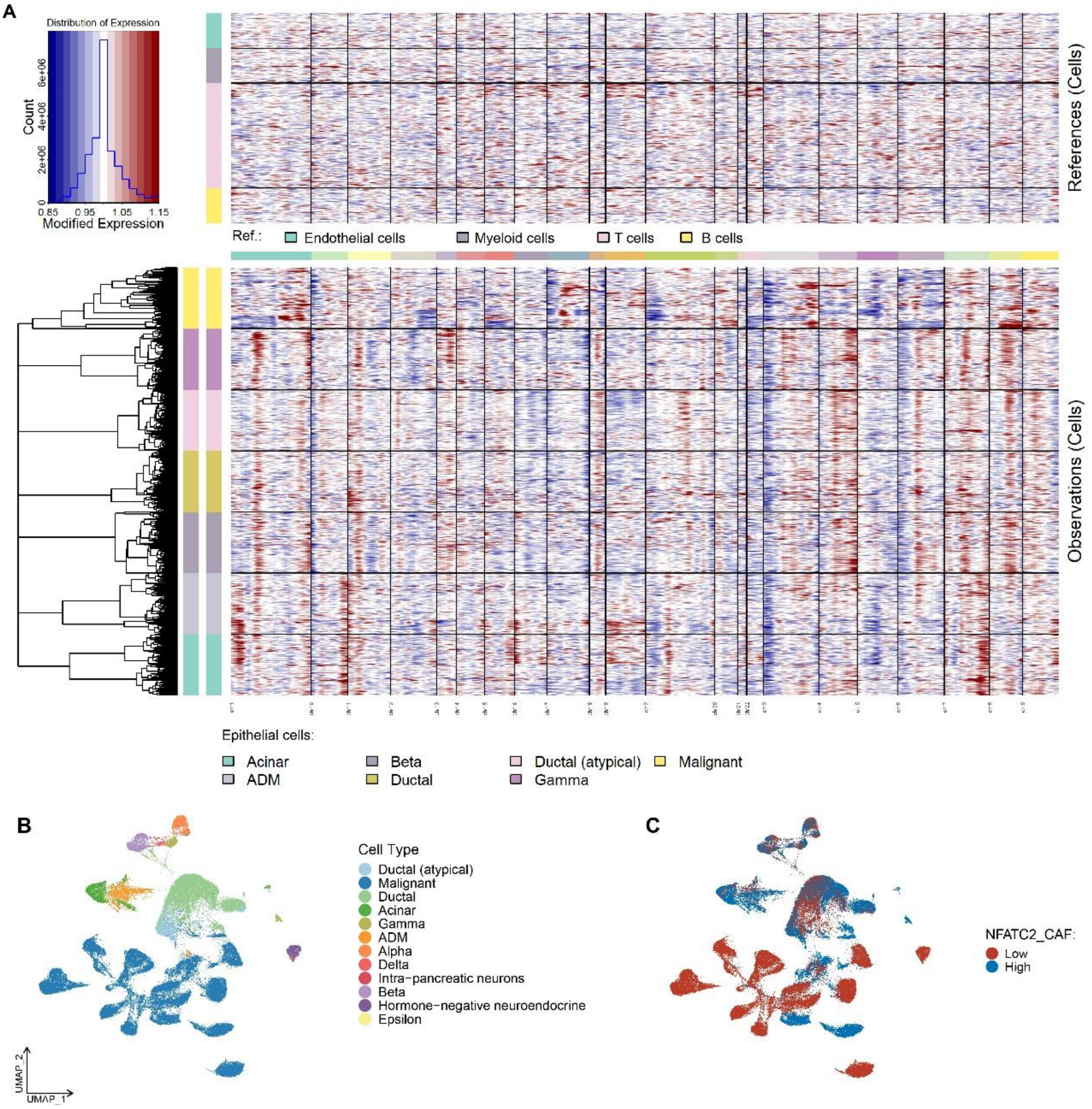


**Supplementary Figure 5. Copy number variation (CNV) analysis to distinguish malignant cells from normal epithelial cells. Related to Figure 5.**

(A) InferCNV heatmap showing chromosomal copy number alterations across epithelial cell populations. Top panel: reference normal cells; bottom panel: test epithelial cells with hierarchical clustering. Red indicates gains, blue indicates losses. Cells cluster into distinct groups based on CNV patterns, with malignant cells showing extensive chromosomal aberrations.

(B) UMAP of epithelial cells colored by cell type annotation. Ductal (atypical), malignant, normal, acinar, gamma, ADM (acinar-to-ductal metaplasia), alpha, delta, ductal-pancreatic neurons, beta, hormone-negative neuroendocrine, and epsilon populations identified based on CNV status and marker gene expression.

(C) UMAP of epithelial cells colored by NFATC2+ CAF abundance (high in red, low in blue), demonstrating spatial distribution patterns.

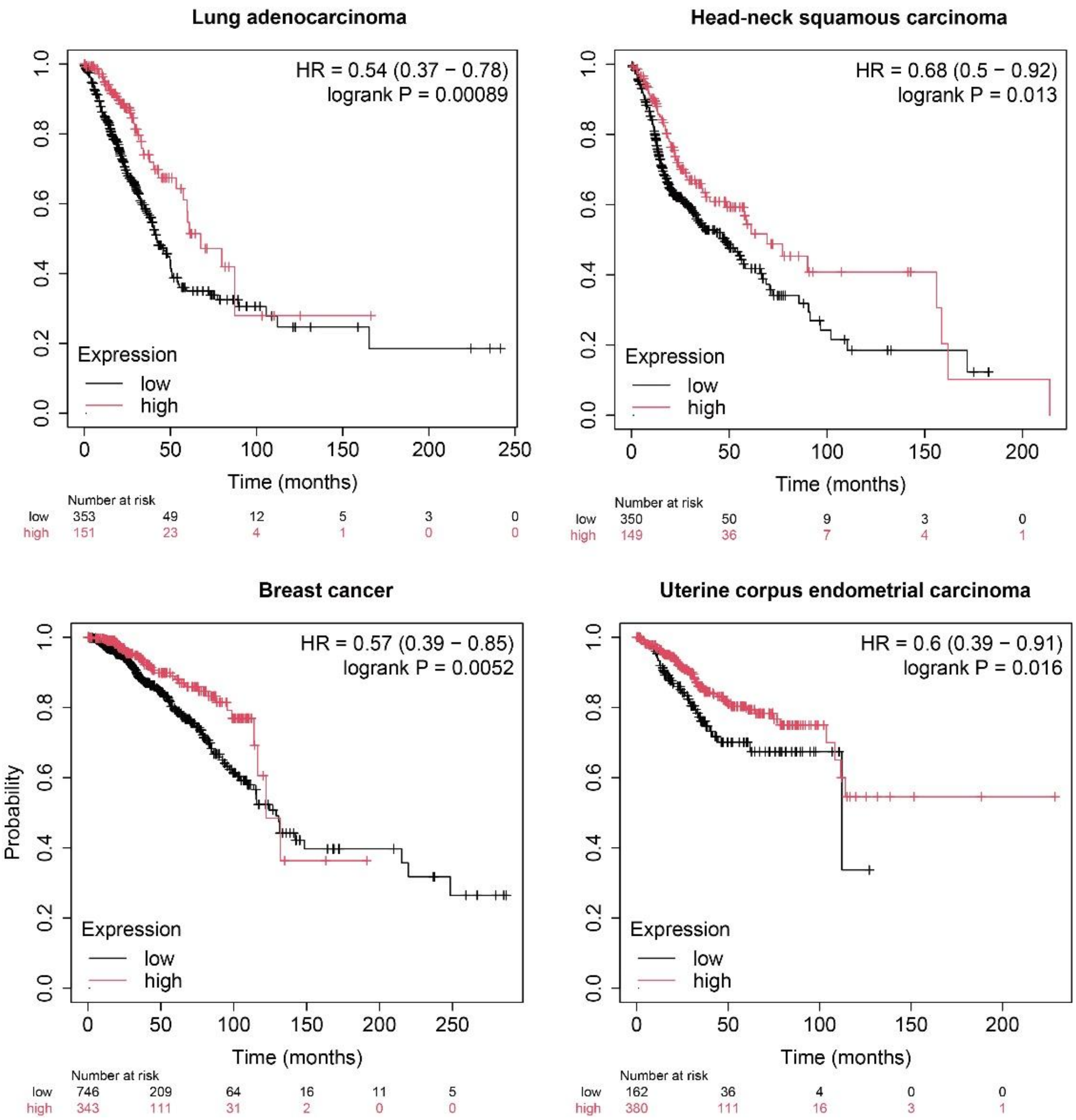


**Supplementary Figure 6. Pan-cancer survival analysis demonstrates prognostic value of NFATC2 expression across multiple tumor types. Related to Figure 6.**

Kaplan-Meier survival curves showing overall patient survival stratified by NFATC2 expression levels (high vs. low) across four different carcinoma types from TCGA datasets. Each panel displays survival probability over time with hazard ratios (HR), 95% confidence intervals, and log-rank p-values. Red lines represent high NFATC2 expression groups, black lines represent low NFATC2 expression groups.
