## Supplementary figures and images for "NFATC2 in pancreatic cancer-associated fibroblasts predicts treatment response and facilitates ERBB-targeted therapies"

### Figure abstract

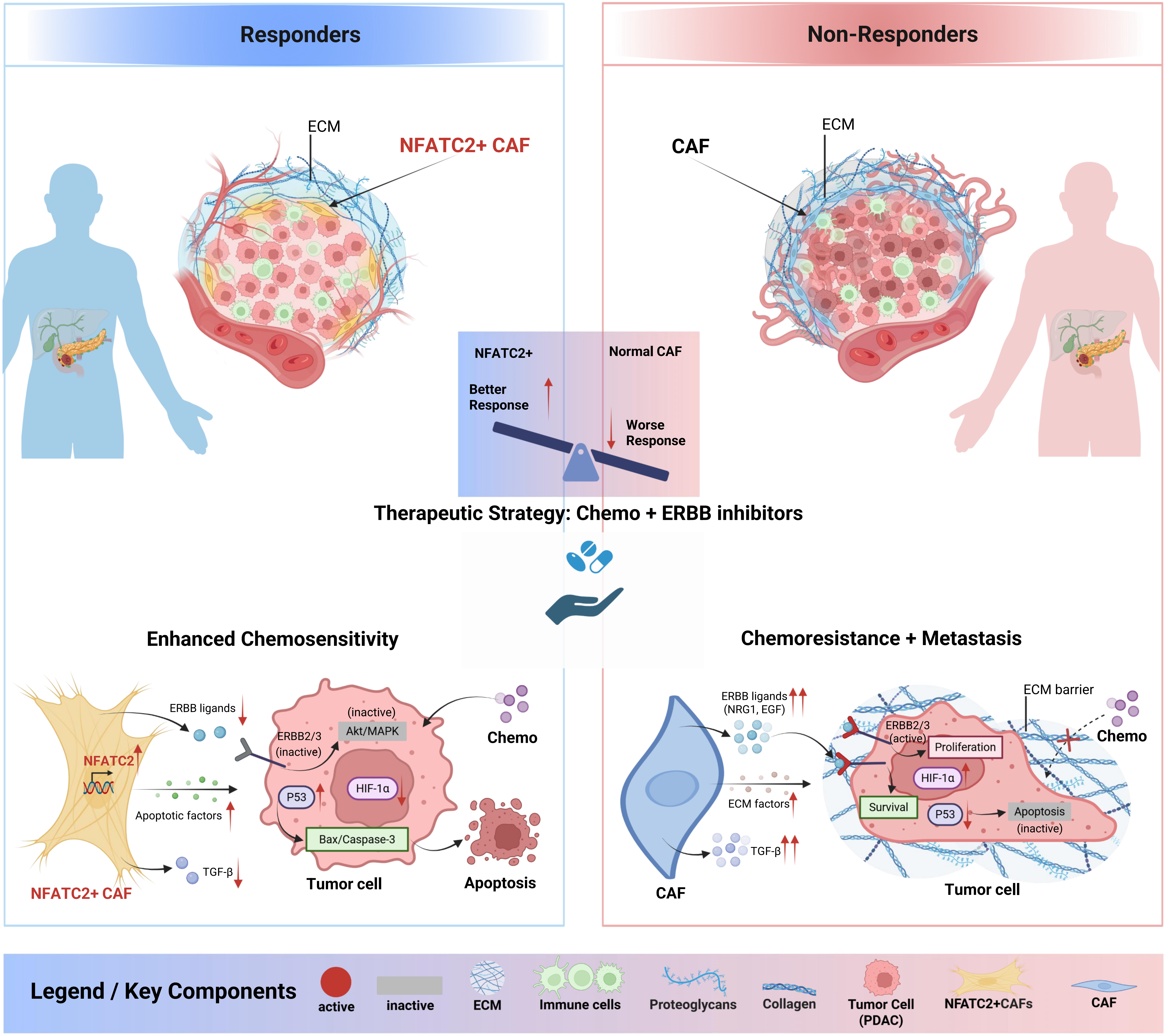
